# On the origin of photoperiod non-responsiveness in barley

**DOI:** 10.1101/2020.07.02.185488

**Authors:** Rajiv Sharma, Salar Shaaf, Kerstin Neumann, Yu Go, Martin Mascher, Michal David, Adnan Al-Yassin, Hakan Özkan, Tom Blake, Sariel Hübner, Nora P. Castañeda-Álvarez, Stefania Grando, Salvatore Ceccarelli, Michael Baum, Andreas Graner, George Coupland, Klaus Pillen, Ehud Weiss, Ian J Mackay, Wayne Powell, Benjamin Kilian

## Abstract

In barley, the transition from the vegetative to reproductive phase is complex and under the control of photoperiodic and temperature conditions. One major gene involved is *PPD-H1*, a *PSEUDO-RESPONSE REGULATOR 7* (*PRR7*) that encodes a component of the circadian clock. Mutation at *PPD-H1* resulted in the photoperiod non-responsive *ppd-H1* alleles that are beneficial under high latitudinal environments as they allow vegetative growth during the long-day summer conditions whereby higher yields are harvested by farmers. Utilizing a diverse GWAS panel of world-wide origin and a genome-wide gene-based set of 50K SNP markers, a strong association of days to heading with the *PPD-H1* gene was detected in multi-location field trials. Re-sequencing of the gene spanning putative causative SNPs, SNP22 (Turner et al. 2005) and SNP48 (Jones et al. 2008), detected recombination between the two, previously reported to be in complete LD. Phenotyping of the recombinants and phylogenetic relationships among haplotypes supported the original conclusion of Turner et al. (2005) that SNP22, present in the CCT domain, is the most likely causative SNP. To infer the origin of non-responsiveness, the *PPD-H1* gene was re-sequenced in a geo-referenced collection of 2057 wild and domesticated barleys and compared with the allelic status of the 6000-year-old barley sample from the Yoram cave in the Masada cliff. A monophyletic and post-domestication origin in the Fertile Crescent was found in contrast to the pre-domestication origin proposed by Jones et al. (2008). We show that the photoperiod non-responsiveness originated from Desert type wild barley in the Southern Levant.

## Introduction

Variation in days to heading affects vegetative growth, the source-sink relation and ultimately determines the yield of the crop. Thus, understanding its genetic architecture is important. Heading date is influenced by various environmental cues. Among them, temperature and photoperiod are the two most important in temperate cereals (McMaster and Moragues 2019). Barley (*Hordeum vulgare* L.), the fourth most important crop, has been traditionally classified as a long-day plant. The species flowers in (early) spring in the Fertile Crescent and during long summer days in north Europe.

Barley is one of the oldest crops and was domesticated in the Fertile Crescent (Harlan and de Wet 1971; Zohary et al. 2013). Wild barley grains have been found in large amounts at the Ohalo II archaeological site on the shore of the Sea of Galilee and were dated to 23,000 years BP (Piperno et al. 2004; Weiss et al. 2004; Snir et al. 2015). This suggests that wild barley was collected from nature long before its domestication. Archaeological data at this site also suggest pre-domestication cultivation and provide evidence that domestication was sometimes not successful (Piperno et al. 2014; Snir et al. 2015). More recent evidence showed that the non-brittle rachis phenotype of domesticated barley originated at least twice, spatially and temporally independent, in the Southern (*btr1*-type) and Northern (*btr2*-type) Levant (Pourkheirandish et al. 2015). The ancient DNA (aDNA) sequencing of 6000-year-old barley grains from the Yoram Cave in the Masada Cliff, is consistent with the origin of barley domestication in the Southern Levant (Mascher et al. 2016).

Migration of the crop outside the Fertile Crescent required adaptation to different environmental conditions (von Bothmer et al. 2003). Present-day barleys are grown in diverse climatic conditions from Scandinavian countries in Northern Europe to the Sub-Saharan desert in Africa and in temperate regions of the Americas and Asia (Dawson et al. 2015), which makes it an important crop for adaptation under a changing climate in the 21^st^ century.

In barley, the transition from the vegetative to reproductive phase is complex (Comadran et al. 2012). Recent knowledge for key flowering genes pathways involved has been reviewed for cereals by Steffan et al. (2014) and Monteagudo et al. (2019). Two major genes involved in photoperiod response have been identified and characterized in barley: *PHOTOPERIOD-H1, PPD-H1* on chromosome 2H (Turner et al. 2005) and *PHOTOPERIOD-H2, PPD-H2* on chromosome 1H (Kikuchi et al. 2011).

*PPD-H1*, is a *PSEUDO-RESPONSE REGULATOR 7* (*PRR7*) gene that encodes a component of the circadian clock (Turner et al. 2005) promoting flowering in both winter- and spring-sown conditions. Photoperiod non-responsive allele(s) (*ppd-H1*) are beneficial under high latitudinal environments as they allow vegetative growth during the long-day spring and summer growth conditions, whereby higher yields are harvested by farmers.

The gene was cloned by Turner et al. (2005) using a winter x spring mapping population. *PPD-H1* consists of 676 amino acids and two major conserved domains described as pseudo-receiver and a CCT (CONSTANS, CONSTANS-like and TOC1).

The wild-type, dominant and photoperiod responsive allele(s) (*Ppd-H1*) at *PPD-H1* accelerate flowering by upregulating *VRN-H3* (*HvFT1*), which is mediated by the activity of CONSTANS (Turner et al. 2005; Campoli et al. 2012). Mutation of *Ppd-H1* resulted in the recessive and photoperiod non-responsive *ppd-H1* allele (Takahashi et al. 1963; Turner et al. 2005).

Two diagnostic single nucleotide polymorphisms (SNPs) differentiating between sensitivity (photoperiod responsive) and insensitivity (photoperiod non-responsive) to long days have been published: Turner et al. (2005) provided strong evidence for a causative SNP in the CCT domain (SNP22), while Jones et al. (2008) described a SNP in exon 6 (SNP48). Re-sequencing different sets of wild and cultivated barleys gave contradictory results. In Turner et al. (2005) and Cockram et al. (2007), the photoperiod non-responsive allele (*ppd-H1*) was not found in wild barley. Thus, the authors concluded that a natural mutation occurred *post-domestication* during the spread of barley cultivation in Europe. Subsequently, Jones et al. (2008) found three non-responsive haplotypes within *wild* barleys from Israel and four non-responsive haplotypes within *wild* barleys from Iran concluding that (I) the non-responsive phenotype of European landraces originated in wild barley from Iran; and (II) that the impact of wild barley from Iran was significant for the domestication history of European barley. However, these results were based on a relatively small sample of potentially admixed wild barleys of genebank origins (Jakob et al. 2014). These non-responsive *wild* barleys were from *east of the Fertile Crescent*, and earlier studies had reported they contributed little to present day European barleys (Kilian et al. 2006; Morrell et al 2007).

In this paper we study the origin and domestication history of photoperiod non-responsiveness (*ppd-H1*) in barley. We first performed Genome-Wide Association Studies (GWAS) of days to heading (Hd) scored in multi-location field trials in a diverse panel of genotypes of worldwide origin. We then re-sequenced the potentially causative genomic region at *PPD-H1* in a comprehensive geo-referenced collection of wild and domesticated barley. Additionally, we used relationships among haplotypes, bioclimatic and phenotypic data, and comparison with the allelic status of the ancient 6000 years old barley sample from the Yoram cave.

## Materials and methods

### The Genome-wide association study panel (GWAS panel)

The diverse spring barley association panel consisted of 127 two-rowed and 97 six-rowed barley genotypes of world-wide origin (Haseneyer et al. 2010; Pasam et al. 2012). One-hundred-and nine genotypes originated from Europe, 45 from West Asia and North Africa (WANA), 40 from East Asia and 30 from the Americas (Table S1). The panel has been successfully utilized in other GWAS for agronomic traits, salt tolerance, drought-stress and candidate gene-based re-sequencing studies (e.g. Stracke et al. 2009; Long et al. 2013; Alqudah et al. 2014, Comadran et al. 2012; Abdel-Ghani et al. 2019).

### Multi-location field trials (GWAS panel)

Days to heading (Hd) was scored in multi-location field trials; four locations in Germany, and one each in USA, Turkey and Syria (Table S2). As the locations were diverse, we treated the four Germany trials as a single location in the GWAS. Hd was scored as days from the date of sowing until 50 percent of the plants had reached growth stage @GS53 (Lancashire et al. 1991). Experimental design and further details are provided in Table S2 and in Supplementary Information online.

### Markers for GWAS analysis

The GWAS panel was genotyped at TraitGenetics GmbH Gatersleben, Germany using the high throughput 50k iSelect SNP chip consisting of 43,461 SNPs (Bayer et al. 2017). All allele calls were manually inspected using GenomeStudio Genotyping Module v2.0.2 (Illumina, San Diego, California). SNPs with more than 10 percent of missing values and > 10% heterozygous calls were excluded. A set of 37,387 SNPs (≥0.05 minor allele frequency) were used for GWAS. SNPs were anchored to the barley reference genome “Morex” assembly (Mascher et al. 2017). Analysis details on the genome-wide association and Site X SNP interaction are provided in Supplementary Information online.

### Geo-referenced Diversity panel for allele mining and phylogenetic analysis

A comprehensive geo-referenced collection (Diversity panel) of 2195 wild and domesticated barley genotypes, from more than one hundred countries, was established to investigate the origin of photoperiod non-responsiveness (*ppd-H1*) in barley. This collection comprises a targeted selection of genotypes described in several publications (Badr et al. 2000; Kilian et al. 2006; Morrell et al. 2007; Jones et al. 2008; Comadran et al. 2012; Pasam et al. 2012, 2014; Tondelli et al. 2013; Jakob et al. 2014; Pourkheirandish et al. 2015; Russell et al. 2016; Xu et al. 2018; Bustos-Korts et al. 2019), extended by newly collected wild barleys i.e. from Israel and Turkey. All germplasm materials were single seed descended (SSD) for at least two generations and spikes were carefully isolated. Based on morphological and taxonomical characterization under field conditions in Germany, 138 samples were not considered for allele mining (Table S3).

For re-sequencing, we considered in total 2057 genotypes (Fig. S4) comprising (i) 942 wild barleys (*Hordeum vulgare* L. ssp. *spontaneum* (C. Koch) Thell., *H. spontaneum*) representing the immediate progenitor of domesticated barleys; (ii) 1110 domesticated genotypes (*Hordeum vulgare* L. ssp. *vulgare*) including 433 landraces from 58 countries, 673 cultivars from 54 countries and 4 others); and (iii) five feral *H. agriocrithon*. (Table S4); Supplementary Information online).

### DNA Amplification and re-sequencing at *PPD-H1*

Genomic DNA was isolated from single leaves of 2057 SSD-derived genotypes with the Qiagen DNeasy Plant Mini Kit (Qiagen, Hilden, Germany), according to the manufacturer’s instructions. The GWAS panel was re-sequenced using two primer combinations. A 1367 bp fragment was considered for analysis. The Diversity panel was re-sequenced using one primer combination only. After trimming, a fragment of 898 bp was considered for multiple sequence alignments. All details are provided in Supplementary Information online.

### Sequence analysis at *PPD-H1*

PCR products were purified by NucleoFast 96 PCR plates (Macherey-Nagel, Germany) and were sequenced directly on both DNA strands on Applied Biosystems (Weiterstadt, Germany) ABI Prism 3730xL sequencer using BigDye terminators. DNA sequences were processed with ABI DNA Sequencing Analysis Software 5.2 and later manually edited by BioEdit 7.2.5 (Hall 1999). Multiple sequence alignments were generated using the *PPD-H1* sequence of cultivar Igri as reference (AY970701, Turner et al. 2005).

Haplotypes were defined using DnaSP v. 5.10.01 (Librado and Rozas 2009). Singleton haplotypes were confirmed by three independent amplifications and re-sequencing.

Sequence diversity statistics was calculated using DnaSP. Diversity-loss, for the total number of sites (Lπ_Total_), and for silent sites (Lπ_silent_) was calculated by: L_π_ = 1-(π_domesticated_/π_wild_) (Tenaillon et al 2004).

Median-Joining (MJ) networks (Bandelt et al. 1999) were constructed using DNA Alignment 1.3.3.2 and Network 5.0.0.1 (Fluxus Technology Ltd., Clare, Suffolk, UK). SplitsTree4 version 4.15.1 was used to generate a NeighborNet planar graph of haplotypes based on the uncorrected P distances (Hudson and Bryant 2006).

### Geographical distribution maps

All maps were prepared using QGIS (https://qgis.org), a free and open source Geographic Information System (GIS) software. All details are provided in Supplementary Information online.

### Experiments to characterize photoperiod responsive and non-responsive genotypes

#### Hd of the GWAS panel under controlled long and short-day conditions

To confirm that haplotype H10 is truly photoperiod responsive, the phenotypic response to daylength (in days to heading) was tested for 41 selected genotypes including (i) 33 photoperiod responsive and non-responsive genotypes of the GWAS panel, and (ii) eight wild and one domesticated barley from the Diversity panel harboring mainly haplotype H10. The tested panel comprised a total of 10 different haplotypes of *PPD-H1* (Table S4; Supplementary Information online.

#### Contrasting the development of photoperiod responsive and non-responsive genotypes

Data published by Alqudah et al. (2014) was explored for this analysis wherein the panel was phenotyped at four developmental stages under inductive long day conditions in the greenhouse. Thermal time was measured as Growing Degree Days (GDD) from sowing to awn-primordium, tipping, heading and anther extrusion stages. The 218 genotypes and their haplotypes considered for this study are shown in Table S4.

#### Heading date under vernalized and non-vernalized long-day field conditions

This experiment was designed to characterize the growth habit and photoperiod responsiveness of genotypes under contrasting vernalization treatments and long-term conditions. In total, 843 wild and domesticated barley genotypes from the Diversity panel were studied in the field at IPK under vernalized and non-vernalized conditions (Table S4; Supplementary Information online.

### Inferring the allelic states of *PPD-H1* and *HvCEN* in the 6000 years old ancient barley sample JK3014 from the Yoram Cave

Ancient barley sequences (Mascher et al. 2016) were retrieved from the short-read archive accession (PRJEB12197) (https://www.ebi.ac.uk/ena/data/view/PRJEB12197). After adaptor removal and merging of overlapping paired-end sequences with leeHom (Renaud et al. 2014), merged reads with a minimum length of 30 bp were mapped to the barley reference genome (version: MorexV2, Monat et al. 2019) using BWA-MEM. Variants (SNP and short indels) were called from uniquely mapped reads (MAPQ >= 20) using BCFtools (Li et al. 2011). The genotype calls were filtered using the following criteria: (i) the minimum reads depth was 1 for homozygous calls, and (ii) the minimum reads depth was 2 in both alleles for heterozygous calls. *PPD-H1* and *HvCEN* sequences (Comadran et al. 2012) were aligned to the MorexV2 assembly BLAST (Altschul et al. 1990). Based on the alignment result, we obtained the relative position in MorexV2 for the previously reported variants of *PPD-H1* (Turner et al. 2005) and *HvCEN*, and retrieved the genotype calls at these sites from our variant calling file.

### Analysis of Environmental Data

In order to investigate and compare the sampling sites of wild, landrace and the ancient barley, 19 GIS-derived historical bioclimatic variables (mean value from 1950-2000) with a spatial resolution of 2.5-min (5 km) were extracted from www.worldclim.org using the R package raster v.3.0-7 (Hijmans et al. 2005). Data was analysed in R (R core team 2018). See Supplementary Information online.

## Results

### Phenotypic data evaluation for the GWAS panel

Large variation was observed for Hd in the diverse GWAS panel across the multi-location field trials (Fig. 1). The trials in Germany showed the largest, and in the USA the smallest variation (Fig. 1; Table S5). The coefficient of variation was high (>4.93) for all trials, reflecting the diversity of the phenotype at each location. Heritability values were moderate to high (0.66 – 0.99; Table S5). We observed significant (P < 0.001, except USA vs. Turkey) and positive correlations between all pairwise Hd phenotypes (Table S6). The geographically closest locations in Syria and Turkey showed the highest correlations followed by Syria and Germany; while correlations with the USA were the smallest.

**Fig. 1:**
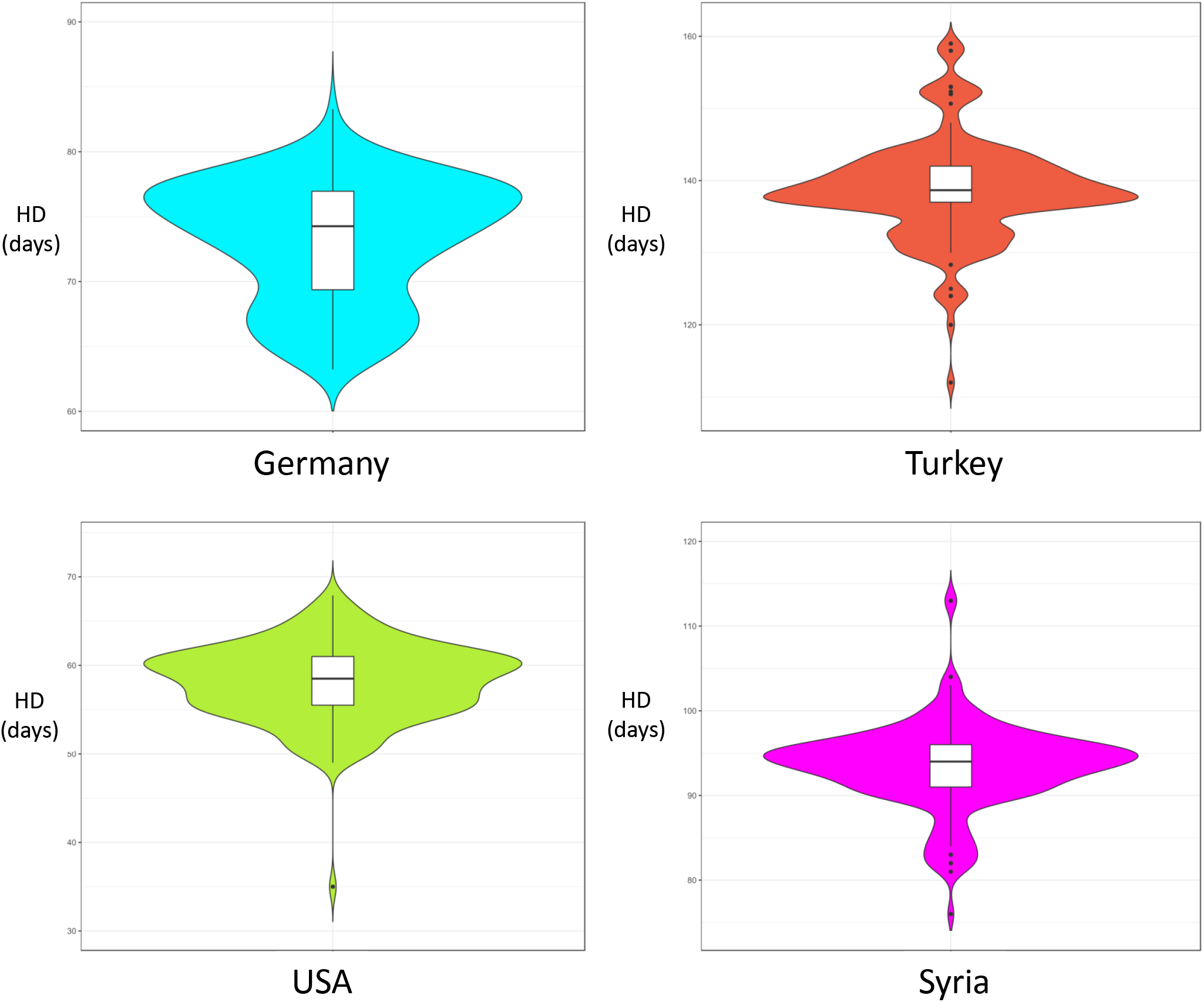
Phenotypic distribution of days to heading (Hd) over four multi-location field trials. Violin plots of Hd (in days) across locations.

### Multi-location heading date associations

A total of 296 SNPs across all sites displayed significant associations (-log_10_P ≥ 4.0) for Hd (Table S7). Most significant regions showed associations with more than one SNP, indicative of the high SNP density and linkage disequilibrium (LD) of the genomic regions detected in the analysis (Fig. 2; Table S7). Despite diverse locations of our trials, major peaks were colocalized near genes like *PPD-H1, HvCEN, VrnH2* and *FT1*. Interestingly, the *PPD-H1* region was not significantly associated for the Montana/USA experiment, which is in agreement with its low pair-wise correlations with the other field sites (Table S6).

**Fig. 2:**
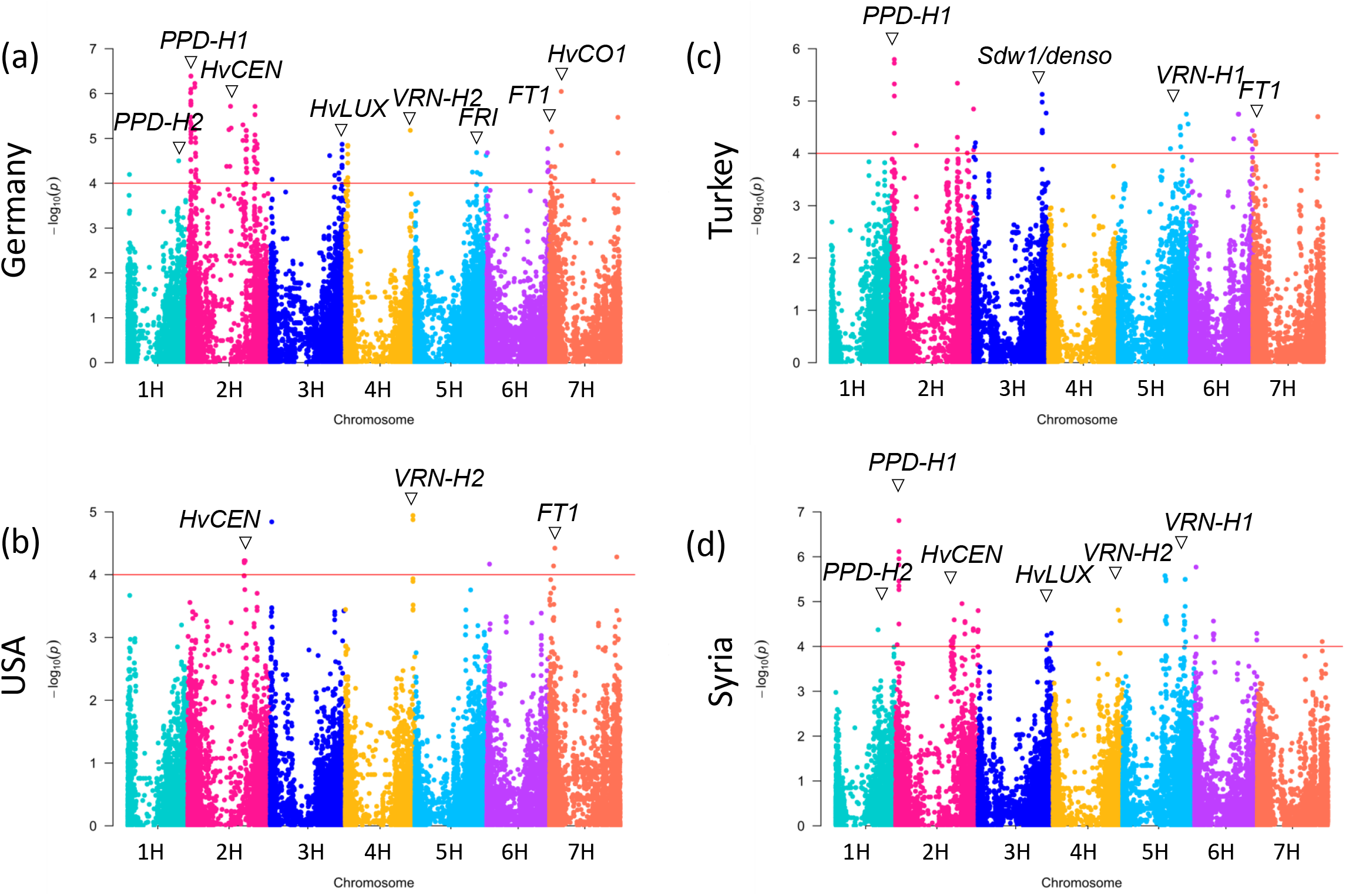
Manhattan plots of days to heading (Hd) from four trial sites are displayed. The horizontal red line shows the significance threshold based on −log_10_P=4.0. (a) Germany; (b) USA; (c) Turkey; (d) Syria. Important co-localized candidate genes are indicated.

Five SNPs at *PPD-H1* showed the highest association for the trials in Germany with significant values up to −log_10_P = 6.39 (Table S7). Association at this region reduces days to heading (Hd) by up to 4.15 days. Further, additional SNPs located on chromosome 2H close to *PPD-H1* were significantly associated indicating the extended LD of the *PpdH1* region (Table S7). SNP48 and SNP22 were analyzed and both were significant with −log_10_P = 6.21 which is similar as the highest associated SNPs on the 50K SNP chip (BOPA2_12_30872 and JHI-Hv50k-2016-73417) (Table S7 and Fig. 3a). Highest associated SNPs were from the intronic region at *PpdH1* (intron 3) and thus less likely to be causative compared to SNP22 and SNP48. Within the 177 accessions for which complete data sets across all locations were available (Table S1), SNP22 and SNP48 were in complete LD (r = 1). However, in the full GWAS panel (224 accessions) these SNPs remained in nearly perfect LD with r = 0.9817 as also observed by Turner et al. (2005) and Jones et al. (2008). SNP22 and SNP48 showed highly significant differences in Hd in Germany compared to Turkey and Syria (Fig. S1). Non-responsive genotypes flowered late compared to responsive types. This could be explained by longer day lengths at higher latitudes in Germany compared to other sites (Table S2).

**Fig. 3:**
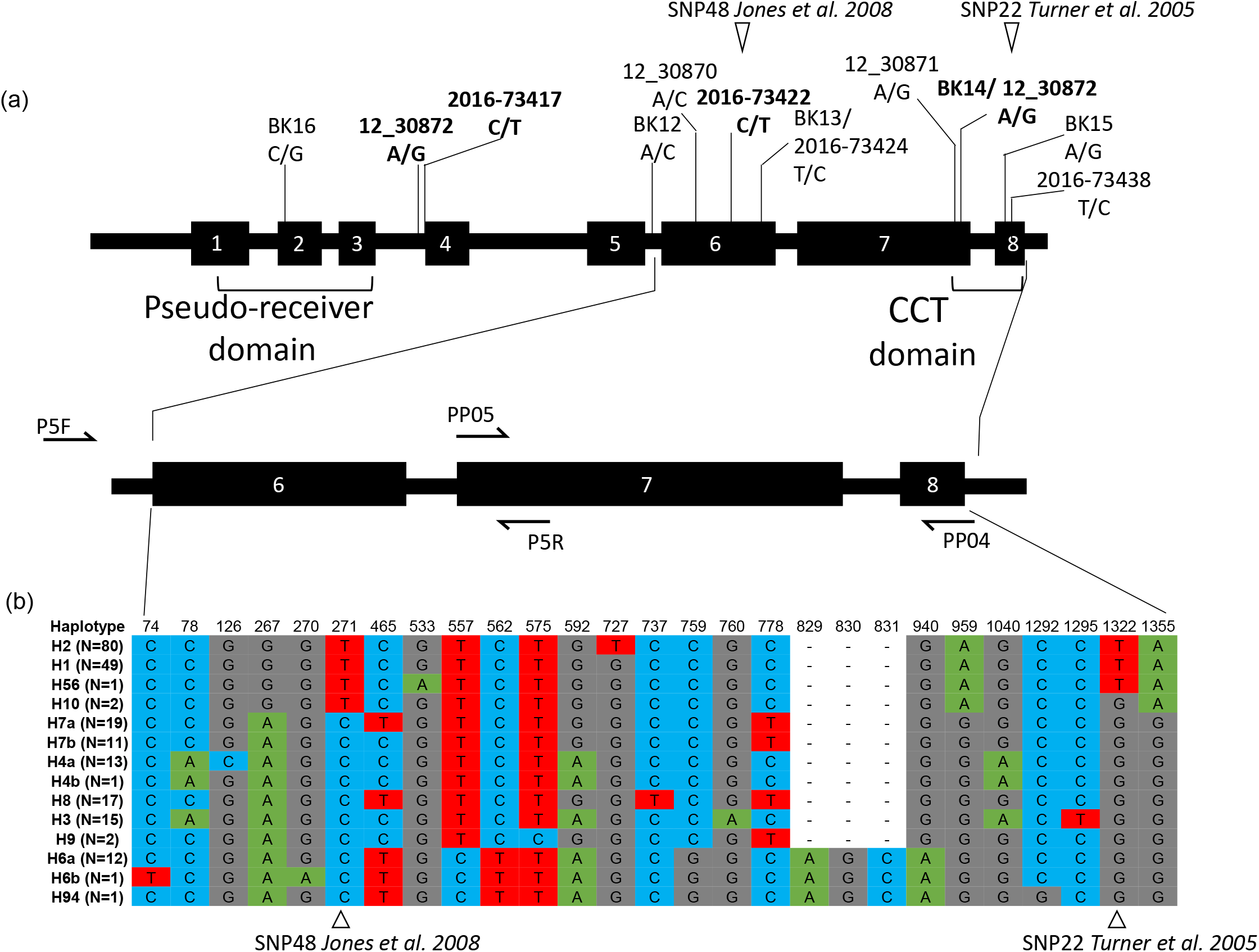
Schematic overview of *PPD-H1* gene structure and SNPs. **(a)** The *PPD-H1* gene consists of 8 exons. Conserved domains are indicated (Pseudo-receiver and CCT). SNP positions are shown. The 50K SNP chip physical positions are given as in Bayer et al. (2017). Details are provided in Table S7; bold=significant SNPs). The locations of SNP22 and SNP48 are indicated. Primer-binding sites are shown by arrows (P5F + P5R; PP05 + PP04). **(b)** SNPs based on re-sequencing at *PPD-H1* in the GWAS panel and corresponding haplotypes (and their frequencies in brackets) are provided.

### Multi-location heading date - SNP interaction

From the significantly associated 258 non-redundant SNPs, 59% (N=153) showed a significant interaction with sites (*P*≤0.05 “p (Site x SNP)”), and 19% (N=51) had a significant interaction with Bonferroni corrected value of 0.05 (Table S8). For 35% (N=54) of significantly interacting SNPs, the relative magnitudes of the interaction and main effects caused the net effect of the SNP on Hd to reverse between at least one pair of sites. For example, the effect on Hd of SNP JHI-Hv50k-2016-73562 near the *PPD-H1* region was predicted as −3.075 days in Germany but +0.739 days in Turkey suggesting contrasting effects of alleles in varying environments (Table 9).

### Haplotype diversity at PPD-H1 in the GWAS panel

The GWAS panel harbored 14 haplotypes (H) (Fig. 3b). Except for genotypes of European origin, which comprised mostly non-responsive haplotypes, no geographical cline in haplotype frequencies was found, suggesting non-responsiveness was selected in European barleys to allow vegetative growth under long-day conditions (Table S4). The most prevalent haplotypes in the panel were haplotype H2 (36%) and H1 (22%). Most European barleys carried haplotype H2 (65%) and haplotype H1 (24%) (Table S10). Barleys from WANA carried in total 11 haplotypes, with intermediate to low frequencies. Frequent haplotypes within American barleys were haplotype H1 (34%) and H8 (25%), while haplotype H6b was predominant in East Asia (23%). Six haplotypes were region-specific (Table S10).

Photoperiod non-responsive haplotypes should carry nucleotide T at both positions, at SNP22 and at SNP48, compared to the cultivar Igri as detected by Turner et al. (2005). Three haplotypes (H1, H2, H56) carried nucleotide T at both positions. Interestingly, haplotype H10 carried T at SNP48 but G at SNP22 and was found in two cultivars (BCC533 and BCC759, from Nepal and India, respectively) (Fig. 3b; Fig. S2; Table S4).

To ascertain whether these two genotypes are photoperiod responsive or non-responsive, both were grown together with 39 further genotypes from the Diversity panel under long and short day conditions. For Hd under long and short-day conditions, repeatability was high with 0.98 and 0.91, respectively. While there was no difference among the haplotypes in short day condition, under long days a clear effect was observed. The seven genotypes carrying H10 were found to be photoperiod responsive (Fig. 4), as haplotype H10 reached heading date significantly earlier than its derived haplotype H1 under long day conditions (p<0.001, 22 days earlier), while under short days, the observed difference of 3.8 days was not significant. This demonstrates that haplotype H10 is photoperiod responsive and concludes SNP22 as the most likely causative SNP. Supporting our findings, both domesticated H10-genotypes of the GWAS panel were early under long day conditions from the tipping until the anther extrusion stage in the study of Alqudah et al. (2014) (Fig. S3). Accordingly, these two genotypes headed also earlier than the non-responsive genotypes of the GWAS panel in field trials in Germany (across four locations and two years), corroborating that haplotype H10 is indeed photoperiod responsive. Therefore, our data strongly suggest that SNP22 located in the CCT domain is the causal SNP.

**Fig. 4:**
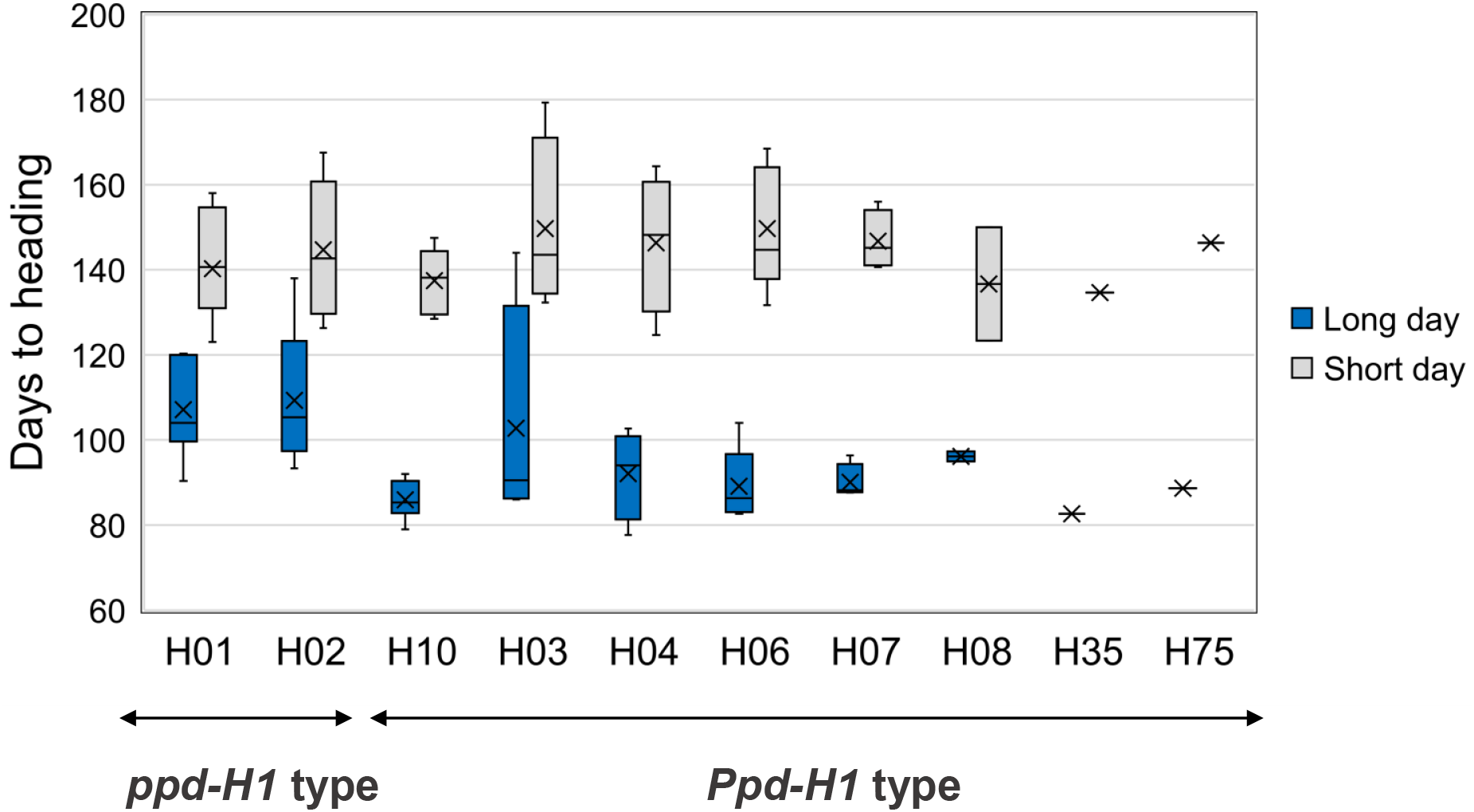
Comparison of phenotypic response (in days to heading at BBCH55) to photoperiod for 41 genotypes grown under long and short-day conditions. Differences in HD under long and short conditions are smaller for non-responsive (*ppd-H1*) genotypes compared to photoperiod responsive (*Ppd-H1*) genotypes. This provides evidence that haplotype H10 containing genotypes are photoperiod responsive. Boxplots are based on phenotypic BLUEs of genotypes.

### Genetic diversity at PPD-H1 within the Diversity panel

We detected ninety haplotypes in 2057 re-sequenced and taxonomically confirmed SSD-derived genotypes (Fig. 5; Figs. S4, S5; Table S13). Higher diversity was found in wild than in domesticated barleys: 71 haplotypes in wild and 27 in domesticated barleys (23 haplotypes in landraces and 17 in cultivars). Eight haplotypes were shared between domesticated and wild barleys. 17 haplotypes were unique to domesticated barley including the non-photoperiod responsive haplotypes H1 and H2. Most haplotypes detected in wild barley had low frequencies. Only haplotypes H6 (40.44%), H4 (14.33%) and H7 (13.69%) were more frequent and were found in 68.46% of the wild barleys (Fig. 5; Table S11). These major haplotypes were not exclusive to wild barley indicating extensive post-domestication utilization.

**Fig. 5:**
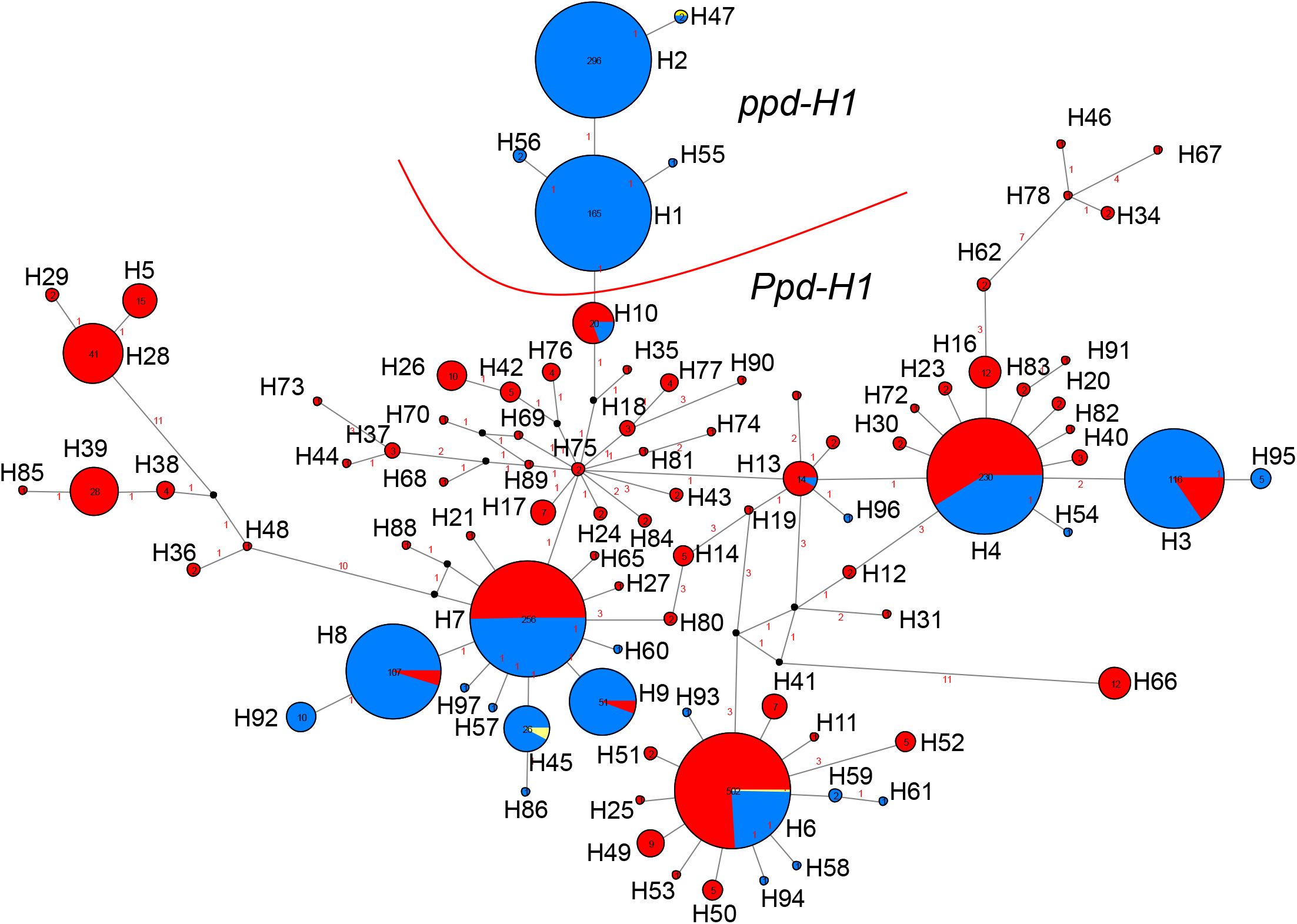
Allele mining at *PPD-H1*. Median-joining network derived from re-sequenced DNA haplotypes of 2057 geo-referenced barley genotypes. 90 haplotypes were found and are represented by arbitrarily given roman numerals. Circle sizes correspond to the frequency of that particular haplotype. Red, haplotype found in wild barley; dark blue, domesticated barley; yellow, *H. vulgare agriocrithon*. Distance in bp between haplotypes is indicated by Arabic numerals and visible at higher magnification. Eight haplotypes were shared among wild and domesticated barleys. Photoperiod non-responsive (*ppd-H1*) haplotypes (H1, H2, H47, H55 and H56) are indicated. Black dots indicate median vectors. Numbers within circles correspond to the number of individuals carrying that haplotype.

Wild barleys from Israel possessed the highest genetic diversity (47 haplotypes), followed by Turkey (19 haplotypes). Several haplotypes were region-specific. Sixty-three haplotypes were unique to wild barley and not exploited in the domesticated barleys, which might harbor a key to local environment adaptation. Interestingly, several of the non-exploited haplotypes clustered together (Fig. 5; Table S11; Supplementary Information online).

Within domesticated barleys, the most frequent haplotypes were H1 (15%) and H2 (27%). These two haplotypes were exclusively found in domesticated barley. Haplotype H1 and its four derived haplotypes (H55, H56, H2, H47) all carried “T” at SNP22 and were considered as photoperiod non-responsive (Fig. 5; Tables S4, 13).

Additionally, we detected 23 haplotypes in 128 genotypes that were classified previously based on their passport data as wild barleys (Table S3; Jones et al. 2008). But careful inspection of them by growing them in the field and studying plant morphology and gene sequence data, we found that they either carrying introgressions from domesticated barley or are seed contaminants from *ex situ* maintenance and seed sharing among genebanks as also reported in Jakob et al. (2014). Thus, we classify them as admixtures and were excluded from the analysis (Supplementary Information online).

Interestingly, in the Diversity panel, haplotypic diversity for wild (H=0.79) and domesticated (H=0.86) barleys was comparable (Table S14). Nucleotide diversity (π) and Waterson’s Theta (θ) were lower for domesticated barley than for wild barley. Differences in nucleotide diversity between wild and domesticated were larger at silent sites (π silent). Within the Diversity panel, 59 segregating sites were found for wild barley that were monomorphic in domesticated barley.

By contrast, domesticated barleys carried 13 segregating sites that were monomorphic in wild material. Tajima’s *D* was negative (-1.55) (P > 0.10) for wild barleys but positive for domesticated barley (0.32, P > 0.10). Only, non-synonymous sites showed significant negative Tajima *D* values (-1.84, P < 0.05) in wild compared to domesticated barleys (0.93, P > 0.10) indicating the presence of rare alleles in wild compared to domesticated barleys. Further, higher Tajima’s *D* values (non-Syn/Syn ratio) were observed for wild (3.55) compared to domesticated (3.01) barley. By comparing wild and domesticated barleys, a 32% loss of diversity (Lπ_silent_) at the silent sites and 9% (Lπ_Total_) at the total number of segregating sites was observed, indicating a relatively moderate diversity loss post-domestication.

### Phylogenetic relationships and geographic distribution of photoperiod responsive and non-responsive haplotypes

Phylogenetic relationships between 90 haplotypes were visualized using a MJ-network (Fig. 5). The central position in the MJ network is occupied by haplotype H75 carried by two wild barley genotypes from Israel (Shilat) located on the western slopes of the Judea mountain ridge. This region was previously identified as a hybrid zone between the Desert and Coast wild barley ecotypes (Hübner et al. 2009, 2013). From here, major geographical groups of haplotypes are visible:

#### Geographical group 1

from H75 to H10 to all photoperiod non-responsive haplotypes: photoperiod non-responsive haplotypes (H1, H55, H56, H2, H47) were exclusively found in domesticated barley (and one *H. agriocrithon*), clustered together and originated from the photoperiod responsive haplotype H10.

Interestingly, haplotype H10 was found in 16 wild barleys (13x Israel, 3x Iran), two landraces (1x Turkey, 1x Nepal) and two cultivars (1x India, 1x Japan) (Table S4, Fig. S6). Among the wild barleys from Israel, 12 were collected in the south and east of Israel where the climate is dry and warm. They were characterized as Desert ecotype and showed early flowering (Hübner et al. 2013). One genotype was collected in the hybrid zone between Desert and Northern ecotypes (FT138, Moledet). Most importantly, among the wild barleys from Israel were eight genotypes recently collected by Hübner et al. (2009) that provides convincing evidence that haplotype H10 exists in wild-stands in nature and that it is associated with the early flowering Desert ecotype. The remaining four genotypes from Israel were collected in the 1960s and 1970s. Also, these showed truly wild characteristics and were collected approximately from the same locations as the eight genotypes by Hübner et al. (2009) (Fig. 7; Table S4). In a recent study wild barleys collected apart 28 years from Isarel were shown earlier flowering phenotype another sign of natural selection iminged on wild stands of present day due to global warming (Qian et al. 2019).

Also among the wild genotypes harboring haplotype H10 were three genotypes from Iran, collected between 1952-1958 in the province of Khuzestan (Table S4).

Haplotype H10 was not found in Turkey despite extensively sampling of the entire distribution area of the species in the country (N=362). The disjunctive distribution of haplotype H10 suggests similar environmental conditions at the respective collection sites.

The non-responsive haplotype H1 was found in domesticated genotypes mostly from Central Europe (Table S4). Haplotype H2, was mainly found in genotypes originating from North-West Europe (Fig. S4). Thus, geographical distribution differences of non-responsive and responsive haplotypes were observed (Figs. S4, S7). Interestingly, the non-responsive haplotypes H1 and H2 were also found in domesticated barley from the Fertile Crescent including in Bedouin landraces, newly collected by Hübner et al. (2009) (Supplementary Information online).

#### Geographical group 2

H75 -> H7 -> H8 -> H92: Haplotype H7 was found in 256 genotypes of almost equal frequency in wild (13.7%) and domesticated (11.4%) barleys. Among the wilds, haplotype H7 was found in 33.3% of wild barley from Israel and 0.6% genotypes from Gaziantep/Turkey. Also, 88 landraces predominantly collected in North Africa and the Near East harbored this haplotype. Haplotype H8 was found in 0.5% of wild barley (4x ISR, 1x CYP) but in 9.2 % of domesticated barley, representing mostly landraces collected in the eastern Mediterranean. Haplotype H92 was mainly found in landraces from Algeria.

#### Geographical group 3

H75 -> H13 -> H4 -> H3 -> H95: Haplotype H13 was detected in 13 wild and one landrace barley from Chad. Haplotype H4 represents a major haplotype and was detected in 230 genotypes (14.33% of wild and 8.55% of domesticated). Wild barleys were collected in Greece (N=1), Israel (N=54, including the Desert type barley FT143), Jordan (N=6), Lebanon (N=4), Syria (N=6) and Turkey (N=64). The remaining were 39 landraces mainly from the Fertile Crescent but also from Libya (N=3), and 56 cultivars mainly from Turkey. All wild barleys harboring haplotype H3 (N=18) were collected west of Gaziantep in Turkey (Supplementary Information online).

### The 6000 years old domesticated barley sample from the Yoram cave was photoperiod responsive

We determined the allelic states of *PPD-H1* and *HvCEN* in the aDNA sample JK3014 extracted from barley grains found in the Judean desert and dated to 6000 years BP (Mascher et al. 2016). After aligning previously published ancient DNA sequences (sample JK3014) to the current barley reference genome sequence (MorexV2, Monat et al. 2019), the genotypes of putative causal variants in *PPD-H1* and *HvCEN* were determined (Table S15). The ancient barley carried the ancestral alleles at *PPD-H1* (photoperiod responsive, G at SNP22) and *HvCEN* (‘early’-flowering, C [proline], at position 531 of Comadran et al. 2012). Multiple sequence alignment analysis concluded that JK3014 carried the following haplotypes; H4 at *PPD-H1* and IV at *HvCEN*. This haplotype combination has not been found in the comprehensive Diversity panel consisting of 2057 genotypes collected in the last 150 years. However, H4 is one of the major haplotypes of *PPD-H1* and shared by wild and domesticated barleys. Interestingly, *HvCEN* haplotype IV was not found in wild barley, it is derived from haplotype II (Comadran et al. 2012). Haplotype IV was found in 21 domesticated barleys mainly from Ethiopia (N=10) but also from Turkey (N=2), Syria (N=1) and Yemen (N=1) (Table S4).

### Analysis of environmental data shed more light on the region of origin ofphotoperiod non-responsiveness

To characterize the collection sites, bioclimate variables were clustered by employing PCA. The precipitation-related variables i.e. Bio14, Bio17, Bio12 and Bio18, were separated from the temperature-related variables (Bio1, Bio5, Bio6, Bio9, Bio10, Bio11) with PC1. With PC2, Bio4, Bio7 were separated from Bio8, Bio13 and Bio16 (Fig. S9; Fig. 6). The PC1 separated mainly collection sites of non-responsive barley carrying the haplotypes H1 and H2 from responsive barleys (other haplotypes). Most collection sites of non-responsive genotypes had higher values of the precipitation-related variables and lower values of the temperature variables (Fig. S10). Interestingly, the PC2 separated non-responsive barleys containing haplotype 2, into two groups (Fig. 6a). One group consisted mainly of landraces from Ethiopia that showed strong correlations with precipitation-related variables Bio13 and Bio16. The second group comprised 40 landraces from 19 countries including Egypt and Israel (Table S16). Especially based on the temperature-related variables (Bio1, Bio2, Bio5, Bio9 and Bio10), the collection sites of H10 containing wild barley from Israel were more closely related to the collection sites of non-responsive H1 containing domesticated barley than the collection sites of H10 wild barley from Iran (Fig. 6a-c; Table S16).

**Fig. 6:**
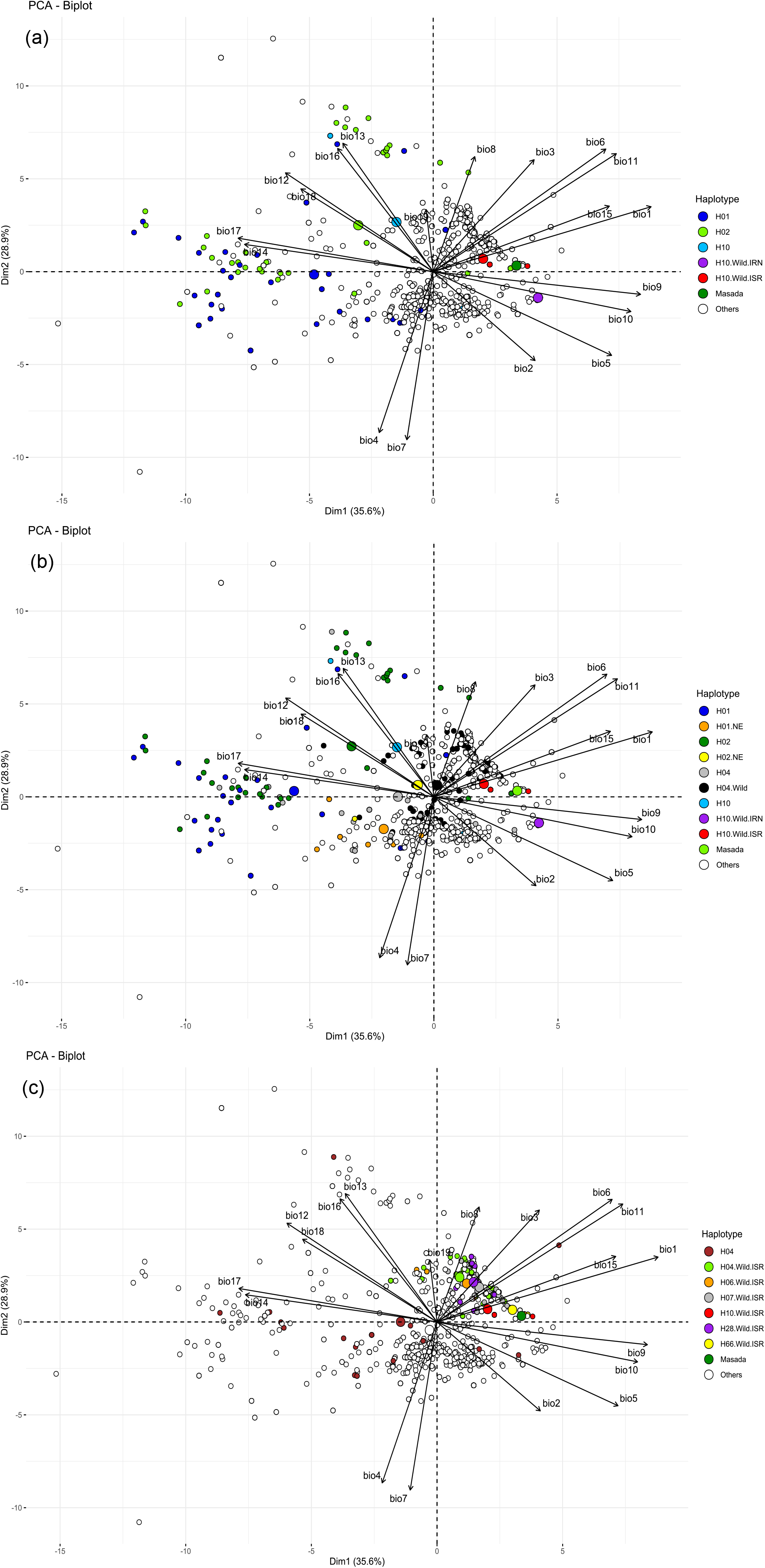
Principal component analysis (PCA) biplot of bioclimatic variables for collection sites of 1375 genotypes (942 wild and 432 landrace barleys, and the Masada sample). **a)** non-responsive (H1, H2) and responsive groups are separated. Two groups of H2 collection sites are visible. H10.Wild.Israel: wild barley from Israel containing haplotype; H10.Wild.Iran: wild barley from Iran containing haplotype 10. **b)** collection site characteristics of non-responsive barley (.NE – landraces from the Near East) and H4 containing landrace barley are highlighted. **c)** Zooming in on the characteristics of the collection sites of extant wild barley from Israel compared to the collection sites of the Masada sample and landraces containing H4. Wild barley yieldcontaining six haplotypes are indicated. Large circles indicate the median PC projection for each haplotype. See Tables S16, S17 and Figure S9 for more information.

In general, landraces containing H1 from the Near East were collected in regions with higher precipitation (Bio12, Bio14, Bio17) than the H10 wild barley (Fig. 6b). H10 containing landraces were collected from contrasting environments. Collection sites of Desert type wild barley from the Southern Levant were positively associated with Bio1, Bio6, Bio11, and Bio15 (Fig. 6c; Supplementary Information online).

Current environmental conditions at the collection sites of extant wild and landrace barley were compared with the collection site of the ancient Masada barley (JK3014) on the basis of pairwise Euclidean distances and bioclimatic variables. The following results were obtained (Fig. 7; Table S18): (i) the environmentally overall most similar collection site of all extant wild barley to Masada was the collection site of the wild barley population B1K-12 (Israel, Kidron stream), where all wild barley was described as Desert types (Hübner et al. 2009, 2013). The *PPD-H1* haplotypes found in this population were H6, H7, H28 and H66; and the haplotype combinations for *PPD-H1* and *HvCEN* were H66HII (FT045) and H7HII (FT047), which are photoperiod responsive and ‘early’ flowering. Most interestingly, also the closest wild barley to *btr1* was collected here (FT643, Pourkheirandish et al. (2015); and (ii) The environmentally second closest collection site, which is also the closest H10 collection site, to Masada is Almog, where FT639 (Desert type) was collected. From this we concluded that the *HvCEN* haplotype II (progenitor haplotype of IV), but also the *PPD-H1* haplotype 10 were found very close to the Masada cliff. The ecologically closest landrace collection site (with exact details in the passport data) to Masada was found in Egypt supporting Mascher et al. (2016) (Table S18; Supplementary Information online)

**Fig. 7:**
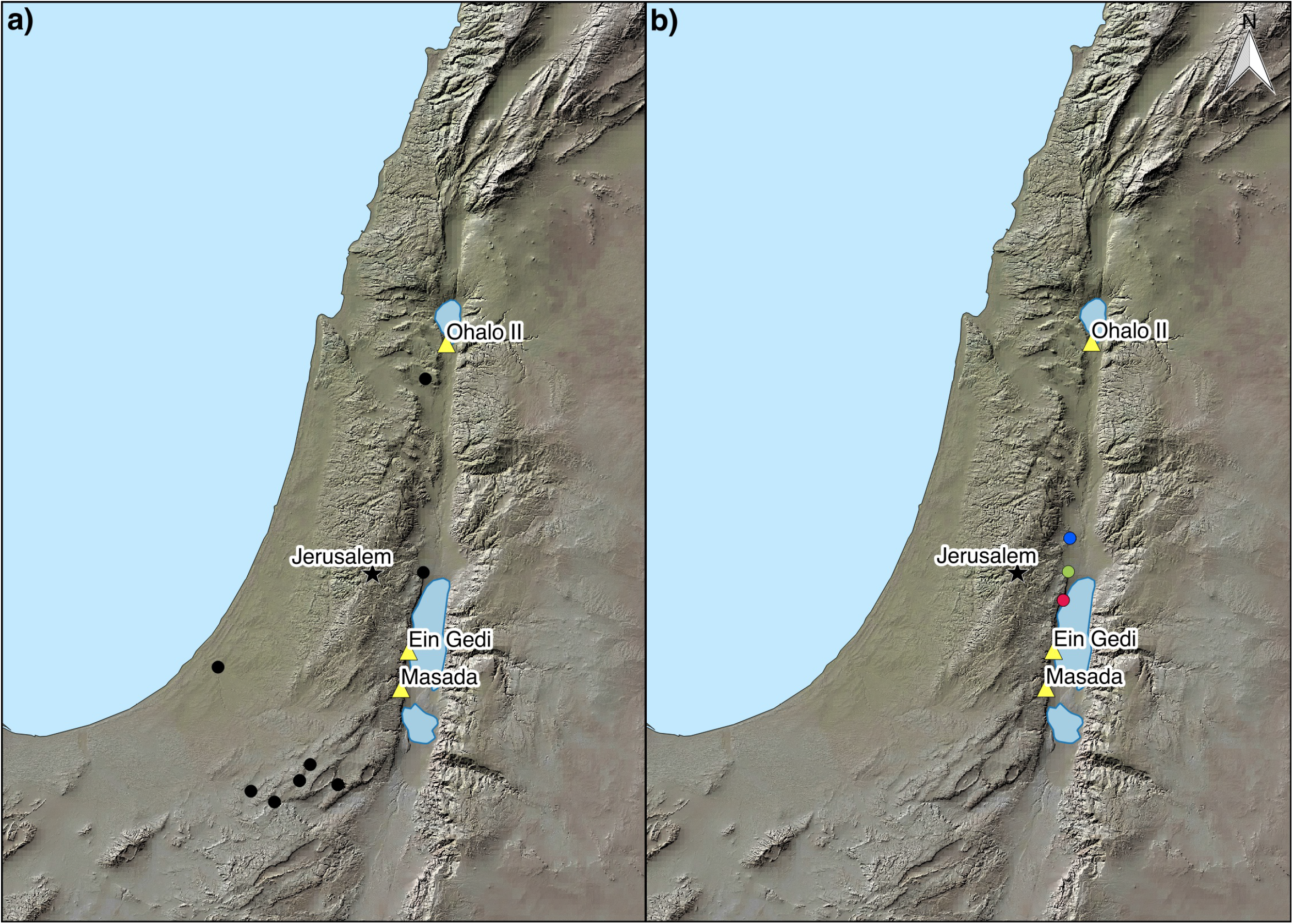
Distribution of *PPD-H1* haplotype 10 in wild barley from Israel, relevant excavation sites, and the environmentally closest collections sites to Masada. a) haplotype 10 collection sites are indicated by black dots. b) **Red dot** – Kidron stream, the environmentally overall most similar collection site of extant wild barley to Masada. FT643 the closest wild barley to *btr1* based on Pourkheirandish et al. (2015) was collected here; **Green dot** – Almog, the environmentally second closest and also the closest *PPD-H1* haplotype 10 collection site to Masada; **Blue dot** – Neomi, the environmentally third closest collection site to Masada. FT013, FT015 and FT016 from this collection site were also among the closest wild barleys to *btr1*. Star – location of Jerusalem; yellow triangles – excavation sites (Ohalo II, Masada, Ein Gedi). See Tables S4, S18).

### Vernalization requirement and phenotypic performance of genotypes containing haplotype H10 under long-day field conditions

Under vernalized and non-vernalized conditions, heading date of 843 genotypes of wild and domesticated barley was investigated to determine their vernalization requirement and to characterize key agronomic traits in the vernalized treatment. For all traits, the repeatability of data was high and ranged from 0.86 to 0.98.

In the non-vernalized treatment, 582 genotypes were heading, while the remaining 261 genotypes (30%) were not flowering and therefore considered winter types. On average, heading date was reached 15 days later under non-vernalized compared to vernalized conditions among the set of 582 genotypes. Interestingly, a bimodal distribution of Hd differences was observed, indicating two phenotypic groups - spring (flowering without vernalization) and facultative growth habit (flowering without vernalization but earlier when vernalized) (Fig. S8a). Of all 843 barley genotypes, 97 wild accessions did not exhibit completely wild characteristics and were excluded from subsequent analysis, leaving 746 accessions for comparison (470 wild and 276 domesticated). A total of 204 accessions were classified as spring types (including 6 wild barleys from Israel) and 305 as facultative, while 237 accessions were classified as winter types.

The genotypes carrying the non-responsive haplotypes H1 and H2 at *PPD-H1* were mostly spring types. In contrast, the photoperiod responsive progenitor haplotype H10 was mainly found in facultative types (wild barley from Israel, N=9; and wild barley from Iran, N=3) but also in two spring types (wild barley FT147 from Israel, landrace FT537 from Turkey) and one winter type (wild barley FT002 from Israel) (Fig. S8b). The genotypes with haplotype H10 showed a short life cycle with the second earliest heading date (1. H66, 2. H10, 3. H26) and the earliest maturity date (1. H10, 2. H66, 3. H26) of all haplotypes under vernalized, long-day field conditions in Germany (Fig. S8c; Table S19), even if only wild barley was considered (Fig. S8d). In addition, the H10-containing genotypes were among the three genotypes with the shortest plant height, narrowest flag leaves, shortest main ear and narrowest main ear width (Table S19). From these data we conclude that plants containing haplotype 10 are well adapted to their local environmental conditions in the Southern Levant or in Khuzestan, and that they are characterized by facultative or even spring growth habit (Supplementary Information online).

## Discussion

We present a GWAS of Hd across multi-location field trials that cover diverse latitudes and longitudes and detected significant associations at the *PPD-H1* genomic region. Using resequencing covering causative SNPs of the *PPD-H1* gene, we found that SNP48 (Jones et al. 2008) is not causal. Further, we re-sequenced the largest SSD and geo-referenced Diversity panel of barley to study *PPD-H1* diversity, phylogeny and domestication history. We arrived at the following five key findings:

### 1. Adaptation of barley depends upon the environment

The onset of Hd, a surrogate measurement of flowering time in crops, takes place in favorable environments. The photoperiod responsive *PPD-H1* gene triggers flowering when the day length increases. In winter barley, the vernalization responsive *Vrn-H2* gene acts as repressor of flowering, although many other gene interactions are also involved. With exposure to cold temperature, the repression gradually reduces and thus *PPD-H1* promotes flowering during the spring period. However, exposure to long photoperiod promotes early flowering in barley which carries the dominant *Ppd-H1* allele, and so reduces the vegetative growth. In higher latitudes (like in Central or Northern Europe), the photoperiod non-responsive *ppd-H1* allele enables vegetative growth during long days (spring season) and thus ensures higher yield. The GWAS analysis of multi-environment trials detected associations from genomic regions corresponding to the major flowering-time genes *Ppd-H2/HvFT3, PPD-H1, HvCEN, HvLUX1, Sdw1/denso, Vrn-H2, Vrn-H1, Vrn-H3/HvFT1* and *HvCO1* (Fig. 2). However, significant effects vary across sites suggesting that for adaptation plants utilize common but variable effects on flowering time, which was also indicated by significant SNP x site interactions. In a recent finding, *PpdH1* and *HvCEN* genes elucidate the Genotype x Environment pattern when grown under spring and winter-sown trials of barley (Bustos-Korts et al. 2019), as we also found in trials across the diverse latitudes (Table S8-S9). Crop plants grown under variable environments experience different photoperiod and therefore the influence of the regulator of Hd varies with the environment (Göransson et al. 2019; Afsharyan et al. 2020). For instance, our lower latitude trials (Turkey, Syria) have less annual variation in photoperiod compared to higher latitude Germany where average day length increases by up to 4 hours during the spring growing season. Furthermore, temperature, humidity and soil type can also influence Hd. The environment specific associations observed in our study indicate that adjustment of Hd depends upon gene expression in each environment. Absence of major significant association at *PPD-H1* in Montana/USA could be due to the adaptation of the genotypes. In Montana, plants must withstand low temperatures and freezing during nighttime, and chilling during daytime until mid-June, followed by a terminal drought. Earlier flowering lines withstand terminal drought environments better. However, in our GWAS panel most domesticated genotypes originated from Europe and were adapted to European conditions and are photoperiod non-responsive and thereby developing more biomass. In the short Montana season, they experienced forced flowering, and this might be the reason that no major associations were found. Notwithstanding this, the small peaks that were observed in the *HvCEN, VrnH2* and *Vrn-H3/ HvFT1* regions and might be important for adaptation under drought conditions such as in Montana.

The region near the *Vrn-H3/ HvFT1* gene showed strong peaks at three sites (Germany, Turkey, USA). This is interesting as it was reported that *HvFT1* is the central regulator where signals are perceived that promote flowering (Nitcher et al. 2013). It is not known why no association was detected in this genomic region in Syria. Either it suggests that different loci in high linkage disequilibrium (LD) but in dispersion contribute to the effects observed in our study or that contrasting alleles within the gene cause these effects which could be supported by observing late heading effects in Syria. Further study is needed to ascertain the effects from the *Vrn-H3/ HvFT1* locus by re-sequencing the region. It has been shown recently that at least four identical copies of the *Vrn-H3/ HvFT1* gene are present in barley varieties that have spring growth habit whereas a single copy is present in most other barley varieties. Therefore, copy number variation causes huge effects on the expression of *Vrn-H3/HvFT1* (Nitcher et al. 2013) and may explain the variation seen in this region. Re-sequencing could shed more light on the observed differences. As shown by Casas et al. (2011), four *HvFT1* haplotypes contributed to differences in flowering time but at the SNP level we could not ascertain the differences precisely.

Our findings confirm Maurer et al. (2015) and Herzig et al. (2018). Under the conditions of the field trials of Halle (Germany) and Dundee (Scotland), the photoperiod responsive wild barley alleles of *PPD-H1* studied in the barley Nested Association Mapping (NAM) population HEB-25 flowered substantially earlier than the non-responsive cultivated barley allele, respectively. In a later study with selected HEB-25 lines segregating for responsive and non-responsive alleles, the early flowering effect of wild barley *Ppd-H1* alleles was verified in Dundee (Scotland), Halle (Germany) and Al-Karak (Jordan), but disappeared under field conditions in Dubai (United Arab Emirates) and Adelaide (Australia) with Hd at day lengths below 13 hours (Wiegmann et al. 2019). The drought conditions in Al-Karak lead to a positive effect of the *Ppd-H1* responsive allele on grain yield.

Further study is needed, but a similar effect as in Al-Karak can be expected in Israel, where the Desert type of wild barley with haplotype 10 was found. In this region, where barleys flower early (before Easter; Passover = Easter is the holyday of the barley harvest), the difference in heading date between haplotypes that respond to the photoperiod, but also between responsive and non-responsive haplotypes could be much less pronounced than in higher latitudes.

Interestingly, the progenitor haplotype H10 of the non-responsive haplotype H1 is associated with the Desert ecotype, which is characterized by early flowering (very early in Almog, around January, when daylength is about 10.5h) to avoid terminal drought. Adaptation to water availability is likely to have led to an orchestra of early alleles at many flowering related loci. As expected, further early haplotypes were found in the Judean Desert. These haplotypes occur together in the Desert type wild barley populations and with different frequencies; e.g. (i) Yeruham: 3x H10, 2x H28; (ii) Neomi: 5x H66; (iii) Shivta: 1x H10, 2x H7, 1x H26, 1x H69; or (iv) Kidron stream: 1x H6, 1x H7, 1x H28, 1x H66 (Table S4). All these haplotypes should have similar effects on Hd in the Southern Levant. Haplotype H10 may not have any advantage over other *PPD-H1* haplotypes in the region. In fact, under field conditions in Germany, all these haplotypes were the earliest for time to maturity, and with the exception of H6, also the earliest in Hd (Table S19). Most genotypes carrying these haplotypes were classified as facultative types (Fig. S8b). The climate in the Judean desert does not favor a strong vernalization requirement, and we conclude that most Desert-type wild barley has a facultative growth habit from which a spring growth habit has evolved. The only six wild barleys with spring growth habit are from Israel: FT050 (H6), FT147 (H10), FT288 (H26), FT301 (H7) and FT019 (H7), which suggests that the spring growth habit originated in the Southern Levant. Support for this hypothesis comes from Saisho et al (2011), who classified seven of 161 wild barley types as spring types, with the majority being facultative. However, no wild barley from Israel was included in their study, and the taxonomic status remains unclear, as certain taxa of hybrid origin, were included.

### 2. SNP22 is the causal basis of the *ppd-H1* mutation

Our GWAS results confirmed that *PPD-H1* is one of the most important genes in regulating flowering and variation as this gene causes natural diversity of barley flowering. In our study the two functional SNPs reported by Turner et al. (2005) and Jones et al. (2008) did not show differences in terms of the significance level. This is in accordance with the observed near perfect LD between these SNPs reported by Turner et al. (2005) and Jones et al. (2008). However, within just 87 genebank accessions, Jones et al. (2008) concluded that the functional SNP22 reported by Turner et al. (2005) is not the causal SNP.

Re-sequencing of the gene space spanning putative causative SNP22 and SNP48, we detected recombination between the two, previously reported to be in complete LD (Fig. 3). In the Diversity panel of 2057 barley accessions we found 20 (0,97%) accessions (16 wild, 4 domesticated) with such recombination (haplotype H10), compared to Jones et al. (2008) who used a landrace panel of mostly European origin and found no recombination between these SNPs. We show that haplotype H10 containing genotypes respond to photoperiod, which leads to early heading under long-day conditions (vernalized) compared to genotypes with non-responsive haplotypes (Fig. 4; Figs. S3, S8). Therefore, SNP22 in the CCT domain of Turner et al. (2005) should be considered as the causal basis of the *ppd-H1* mutation, which is supported by phylogenetic analysis. All photoperiod non-responsive haplotypes (H47, H2, H56, H55 and H1) clustered together in the MJ network, thus suggesting a monophyletic origin of photoperiod non-responsiveness in barley due to the G to T non-synonymous substitution at *SNP22* (Fig. 5). *It is important* to note that in our study no phenotypically wild barley was found that carried a photoperiod non-responsive haplotype (*ppd-H1*). Thus, our data indicate that all extant wild barley is photoperiod responsive (*Ppd-H1*). This is supported by Baloch et al. (2013) studying wild barley from Jordan and Iran.

### 3. Photoperiod non-responsiveness originated from Desert type wild barley in the Southern Levant

In total, sixteen wild barleys harboring H10 were collected from Israel and Iran (and not from the central part of the Fertile Crescent). Based on our analysis of environmental data, we show that wild barley from Israel containing H10 (N=13) grow in more similar habitats to non-responsive barley than H10 wild barley from Iran (N=3). For some of them, the haplotype of a second important flowering time gene, *HvCEN* is known form the study of Comadran et al. (2012). The following *PPD-H1* - *HvCEN* haplotype combinations were found in Israel (3x H10III, 1x H10IX) and Iran (3x H10I). This result supports the origin of European non-responsive barley from Desert type wild barley from the Southern Levant, likely carrying H10III, as haplotype III at *HvCEN* was by far the most frequent haplotype in European non-responsive spring and winter barley (Comadran et al. 2012).

But how can the occurrence of wild barley from Iran with haplotype 10 be explained? We speculate that the province of Khuzestan in southwestern Iran was part of the ancient natural distribution range of the species and that haplotype 10 survived the Last Glacial Maximum (LGM) about 21k years ago in the region (Jakob et al. 2014).

Among the four domesticated barleys harboring the haplotype H10, we found a 2-rowed, naked landrace from Turkey (FT537) that was collected by Jack Harlan in 1948. The allelic status at *HvCEN* is not known. In contrast, the haplotype combination H10I was found in (i) one landrace from Nepal (6-rowed, hulled); (ii) one cultivar from India (6-rowed, hulled); and (iii) one 2-rowed, naked cultivar from Japan (Fig. S6). These four domesticated barleys probably originate from H10I containing wild barleys from Iran. Further study is needed to investigate the contribution of wild barley from Iran to the *btr2* genepool (Pourkheirandish et al. 2015). Remains of non-brittle two-rowed barley dated to the Middle PPNB (10th century BP) were found in sites across the Fertile Crescent: Jericho (Israel, only about 10 km from the haplotype H10 collection site Almog), Tell Aswad (Syria), Jarmo (Iraq) but also Ali Kosh (Iran, Khuzestan province, where H10 wild barley was collected) (Alizadeh (2003); Zohary et al. (2013).

### 4. No severe genetic bottleneck at the *PPD-H1* gene

Often a severe genetic diversity change is observed when comparing wild and domesticated barley populations. Signatures of domestication include reduced genetic variation compared to the wild. We observed 14 haplotypes at *PPD-H1* (1376 bp fragment) within the diverse GWAS panel of world-wide origin (Table S10). Further, in the Diversity panel, we found 90 haplotypes within an 898 bp fragment (Table S11). This number of haplotypes is impressive compared to most re-sequencing studies in barley, but it is strikingly low compared to the 121 haplotypes observed in only 266 accessions (that consisted of only 72 wild barleys) by Jones et al. (2008). Theoretically, this difference could be due to the smaller fragment re-sequenced in our study (898 bp considered) compared to 3508 bp re-sequenced by Jones et al. (2008). Other reasons could be issues with SNP calling and haplotype assignment by Jones et al. (2008).

Comparing nucleotide diversity between wild (n = 942) and domesticated barley (n = 1110), we observed only a 9% loss of diversity at *PPD-H1*. This is also contrary to the findings of Jones et al. (2008) that reported a severe bottleneck and 22.5% loss of diversity within their 72 wild and 194 domesticated barleys. Due to the small sample size, diversity values by Jones et al. (2008) should be noted with caution. We have 61 wild barley accessions in common with Jones et al. (2008) (Table S3; Table S4). Thus, we are convinced that the lower number of 90 haplotypes in our study is robust. More haplotypes described by Jones et al. (2008) probably result from issues with SNP calling and haplotype assignment. Nevertheless, recent studies (Cuesta-Marcos et al. 2010; Russell et al. 2016) reported also lower haplotype numbers at *PPD-H1* even from the larger-sequence lengths and diverse samples including wild barleys than Jones et al. (2008).

In our study we used comparative numbers of wild and domesticated barleys and comparatively smaller loss in genetic diversity was observed (Jakob et al. 2014). One reason could be that several major photoperiod responsive haplotypes are shared among wild and domesticated genotypes and that probably led to the observation of low nucleotide diversity change. One other reason could be that the domesticated group is a mix of spring, facultative and winter types, different row and caryopsis types and was collected from a wide range of environments. Thus, diversity (= expected heterozygosity on random mating) will be pushed higher in the domesticated group as a result of the winter – spring, 2-rowed – 6-rowed and other polymorphisms. There may be a general point that selection that maintains polymorphism within the domesticated group, as here, will also maintain diversity or at least reduce the loss. So, we get a reverse of the usual *pre-post-domestication* pattern with less loss than under neutrality. Nevertheless, overall, we observed fewer haplotypes (N=27) in domesticated barley (landraces N=23; cultivars: N=17) compared to truly wild barley (N=71).

Interestingly, recent investigation in sorghum using sequencing of the archaeological samples of wild and domesticated sorghum of different historical periods revealed that the surge in diversity occurred over time and the formation of a domestication bottleneck is probably a myth (Smith et al. 2019; Brown 2019).We observed within wild and domesticated barleys segregating sites exclusive to either group. Segregating sites exclusive to the wild barley indicates that there are many mutations in wild barleys that were possibly not selected in domesticates. However, mutations exclusive to the domesticated barleys suggest that these mutations probably occurred after the initial domestication and/or outside the natural distribution range. Since wild and domesticated barleys can coexist together in the farmer’s fields, natural gene flow may alter the values of genetic diversity. Such cases are reported to be relatively rare and are unlikely to be important in nature (Abdel-Ghani et al. 2004; Russell et al. 2011; Hübner et al. 2012). However, we believe that the rate of re-introgression of wild barley alleles probably occurred more frequently than previously thought.

To broaden the genetic basis for barley improvement at *PPD-H1*, haplotype information from this study could be considered. Potentially beneficial haplotypes could be introgressed from wild barley into the elite background (Dempewolf et al. 2017: https://www.cwrdiversity.org/project/pre-breeding/). Gene editing will provide another opportunity.

### 5. Photoperiod non-responsiveness most likely originated post domestication and in the Fertile Crescent

Barley domestication history is complex. In recent years publications suggest multiple domestication events that led to the present domesticates (Kilian et al. 2006; Morrell and Clegg 2007; Dai et al. 2012; Zeng et al. 2018; Pourkheirandish et al. 2015). One of the major events that led to the adaptation of barley to wider areas is the evolution of non-responsive barley (*ppd-H1*). Two contrasting hypotheses about the origin and spread of non-responsive barleys were published: (I) Turner et al. (2005) and Cockram et al. (2007) suggested that photoperiod non-responsiveness originated in domesticated barley (*‘post’ domestication, outside of the Fertile Crescent, during the spread of barley cultivation towards Northern Europe*); and (II) Jones et al. (2008) concluded that the non-responsive phenotype originated in wild barley from Iran (*‘pre’ domestication*).

In our study we showed that the origin of photoperiod non-responsive haplotypes was derived from photoperiod responsive haplotype H10, and that the mutation leading to non-responsive types was only found in domesticated barley. Our combined data indicate a monophyletic natural mutation that most likely occurred in a domesticated *btr1Btr2*-type, 2-rowed, facultative barley in the Southern Levant (*‘post domestication in the Fertile Crescent’*).

A likely scenario would be that a domesticated barley with a photoperiod responsive allele at *PPD-H1* e.g. H4, such as the Masada barley, hybridized with a Desert-type, facultative, wild barley harboring haplotype 10, in the Southern Levant, probably where H10 barley still grows today (Fig. 7). The fully fertile F1 hybrid would be a ‘domesticated’ barley harboring, for example, the *PPD-H1* – *HvCEN* haplotype combination H10II or H10III. Their offspring would later receive the natural mutation at SNP22, also in the Southern Levant, probably under irrigated cultivation (if we assume that the non-responsive haplotype H1 would have negative effects on barley in this hot and dry region). In a second scenario, a domesticated *btr1Btr2*-type barley, which already contains H10 would directly receive the mutation at SNP22.

The ancient Masada sample is important in this context. Mascher et al. (2016) concluded that this 6000-year-old sample was a domesticated (*btr1 Btr2*), 2-rowed and hulled barley. We found that it carried the *PPD-H1* – *HvCEN* haplotype combination H4IV and was therefore photoperiod responsive (H4) and *‘early* flowering’ (IV). Interestingly, this haplotype combination was not found in the comprehensive collection of 2057 wild and domesticated barleys. The phylogenetically and phenotypically closest *PPD-H1* – *HvCEN* haplotype combination found in the Diversity panel was H4II, and found in one wild barley from Turkey (Northern Levant, near Gaziantep); in four landraces (3x Libya, 1x Georgia) and in nine cultivars from six countries outside the Fertile Crescent (Table S4). At *HvCEN*, the progenitor haplotype of IV is II, which was found in Israel in 8 wild barleys (also in the environmentally closest collection sites to Masada, Fig. 7); and in one landrace obtained from a market in Jerusalem in 1964. Haplotype 4 was found in Israel in 54 wild barley. Our data suggest that the Masada barley evolved from local populations in the Southern Levant and was not introduced from elsewhere.

The probable source of the Masada barley found in the Yoram Cave is located in the Ein Gedi oasis, about 17 km north of the cave (David 2015; Fig. 7). This is the largest and most important oasis in the Judean Desert with an annual water flow of about 3.5 million cubic meters. There, abundant archaeological remains attest for agricultural irrigation systems of agricultural terraces fed with spring water (Hadas 2012). Although the irrigation systems date to the Roman-Byzantine period (1st-6th century CE), the coexistence of nearby archaeological sites from the Chalcolithic period indicates that the spring was flowing at the same time. In addition, a review of historical sources attests to barley cultivation in this irrigation system during the Roman period (Hadas 2012). Considering this, the barley of the Yoram Cave was probably grown under irrigated conditions in the Ein Gedi oasis. Apparently, this is the only example of irrigated cereal fields in ancient Israel.

Similarly, the origin of non-responsive barley could have occurred under irrigated conditions in the Southern Levant. The mutation probably took place in domesticated barley but also a wild barley with H10 could have been mutated and then hybridized with a domesticated barley, under irrigated conditions in mixed stands.

Few photoperiod non-responsive haplotypes, but with a high frequency and found only in domesticated barley, indicate that they are probably of relatively recent origin. Although the 6000-year-old Masada sample was photoperiod responsive, this does not exclude the possibility that the photoperiod non-responsiveness originated earlier than 6000 years ago or that photoperiod responsive and non-responsive domesticated barley co-existed in the Southern Levant in the past as today (Table S4). Based on von Bothmer at al. (2003), *“Barley cultivation reached Spain ca. 7,000 years BP (Before Present), N Africa and Ethiopia ca. 8,000 years BP and northern Europe ca. 6,000 years BP.”* Further study is needed to shed more light on the age of non-responsive haplotypes.

### Conclusion

We showed that the photoperiod non-responsive adaptation to long day spring conditions in Europe originated from Desert type wild barley (H10) in the Judean desert and involved the selection of one *de novo* mutation (SNP22). Haplotypes H1 and H2 increased in frequency during the spread of civilization out of the Fertile Crescent towards Northern Europe under higher selection pressure. Haplotype H2 probably originated also *de novo* (synonymous substitution) during this range extension. Our data suggest that the spring growth habit evolved from the facultative habit in the Southern Levant. Finally, we conclude that all *btr1*-type barley evolved from Desert type wild barley near the Dead Sea.

## Supplementary Material

Supplementary material mentioned in the text, comprising 19 supplementary tables and 13 supplementary figures are available online.

## Sequence availability

New sequence data from this article are deposited in GenBank Data library under accession numbers provided in Tables S12 and S13 (Supplementary Material online): KF309068-309171.

## Acknowledgements

We thank the following for providing seeds and/ or DNA material: Federal *ex situ* Genbank of Germany in Gatersleben (IPK), Germany; U.S. Department of Agriculture, Agricultural Service (USDA-ARS); International Center for Agricultural Research in Dry Areas (ICARDA); Max Planck Institute for Plant Breeding Research (MPIPZ), Cologne, Germany; Australian Winter Cereals Collection, Tamworth, Australia; Institut für Pflanzenbau und Pflanzenzüchtung, Braunschweig, Germany; Aegean Agricultural Research Institute, Izmir, Turkey; Swedish University of Agricultural Sciences (SLU), Sweden; NPGBI-Karaj, Iran; Seed and Plant Improvement Institute (SPII), Karaj, Iran; the Exbardiv consortium; M. Abbasi, S. Michel, M. Ristow, M. von Korff, R. Fritsch, S. Jakob and F. Salamini. We thank Sigi Effgen, Jutta Schütze (both MPIPZ), Kerstin Wolf, Heike Harms, Birgit Dubsky, Christiane Kehler, Heloise Giraud, Peter Schreiber, Jürgen Marlow, Michael Grau (all IPK) and all gardeners and students involved for excellent technical support. The authors thank B. Steffenson, A. Tondelli, M. von Korff, M. Koorneef and F. Salamini for discussions. This work was supported by the German Research Foundation (DFG) Priority Program SPP1530 (FKZ KI 1465/6-1); the ERA-PG-funded project Exbardiv (http://www.erapg.org), GABI Future GENOBAR (https://www.pflanzenforschung.de), the Leibniz Institute of Plant Genetics and Crop Research (IPK), Gatersleben and the Max Planck Institute for Plant Breeding Research, Cologne.

