## Supplementary Material Notes for "On the origin of photoperiod non-responsiveness in barley"

### **Supplementary Materials and Methods**

### **Multi-location field trials (GWAS panel)**

All field trial Hd data were manually checked and outliers were removed based on Z-test scores >3.5, accounting for two to five observations per trial. Restricted Maximum Likelihood (REML) was used to obtain Best Linear Unbiased Estimates (BLUEs) of mean from each location as implemented in Genstat 18 (Payne 2009). Summary statistics of Hd and heritability were calculated using Genstat. One hundred and seventy-seven accessions with complete data sets across all locations were used for GWAS (Table S1).

### **Markers for GWAS analysis**

Genome wide association studies were performed in TASSEL version 5.2.54 using MLM model (Zhang et al. 2010). A–log_10_ (P-value) = 4 was used as a significance threshold to control false positives across the GWAS. Allelic effects, which in TASSEL are calculated based on alphabetical order, were converted relative to the genotype BCC1391 due to no missing genotypic data. A linear regression model was fitted in R (www.R-project.org/) to estimate Site x SNP interaction, without population structure control from significant SNP markers.

### **Geo-referenced Diversity panel for allele mining and phylogenetic analysis**

Wild barleys from the following countries were considered, from west to east: Libya (N=2), Greece (N=4), Cyprus (N=4), Israel (N=363), Jordan (N=18), Lebanon (N=8), Turkey (N=362), Syria (N=37), Iraq (N=29), Iran (N=91), Afghanistan (N=7), Turkmenistan (N=8), Uzbekistan (N=4), Tajikistan (N=4) and ‘Former Soviet Union’ (N=1).

Based on morphological and taxonomical characterization under field conditions at IPK in Germany, 138 samples were not considered for allele mining (Table S3). Among them were 128 samples formerly assigned as Hordeum spontaneum (based on genebank passport data), but all these samples showed non-wild characteristics - based on morphology (non-brittle/ non-shattering types, no awns, naked caryopsis), or gene sequence data at *Btr1* and *Btr2* loci (Pourkheirandish et al. 2015) and at seven other nuclear loci (Jakob et al. 2014) (Table S3).

### **DNA Amplification and re-sequencing at PPD-H1**

The Primer3 online software (http://bioinfo.ut.ee/primer3-0.4.0/; Untergasser et al. 2012) was used to design two primer pairs for PCR amplification and re-sequencing. Both primer pairs amplified in total 1428 bp of *PPD-H1* (positions 12960 – 14388 of cultivar Morex, AY943294).

The first primer pair (P5F-P5R, Jones et al. 2008) covered SNP48 located in exon 6 and amplified 619 bp of cultivar Morex (positions 12960 to 13579): P5F (forward) 5’- GATGGATTCAAAGGCAAGGA-3’ (12960 to 12976 of AY943294) and P5R (reverse) 5’- CGTTAGAGCCCTGCTTCATC -3’ (13560 to 13579 of AY943294). The second primer pair (PP05-PP04, Jakob et al. 2014) covered the CCT domain encompassing non-synonymous SNP22 of Turner et al. (2005). PP05 (forward) 5’-GTGCAAAGCATAATATCAGTGTCC-3’ (13376 to 13399 of AY943294) and PP04 (reverse) 5’-GGCCAAAGACACAAGAATCAG (14368 to 14388 of AY943294) amplified 1012 bp (positions 13376 to 14388; Turner et al. 2005).

The GWAS panel was re-sequenced using both primer combinations. A 1367 bp fragment was considered for analysis. The Diversity panel was re-sequenced using the second primer combination only (PP05-PP04). After trimming, a fragment of 898 bp was considered for multiple sequence alignments.

### **Geographical distribution maps**

Maps were generated from the geographical coordinates of the original accessions. All coordinates were manually checked and standardized in decimal format. A country mismatch test was performed. This test consisted in mapping the coordinates and comparing whether the records were mapped robustly in the countries of origin reported in the passport data. Records where the country origin did not match with the country where the coordinate was mapped, were discarded. Maps are projected using the World Robinson Projection, a projection that minimizes all types of distortion over the sections of a map. Land cover data with shaded relief and country administrative boundaries maps were made with Natural Earth (naturalearthdata.com). We extracted elevation data for each geo-referenced record using CIAT’s SRTM (<http://srtm.csi.cgiar.org/srtmdata/>) resampled raster data (1 sq. m at the equator).

### **Experiments to characterize photoperiod responsive and non-responsive genotypes**

#### *Hd of the GWAS panel under controlled long and short-day conditions at IPK, Germany*

Two seeds of each genotype were sown per turf tray on August 30^th^ of 2017 and thinned out to one plant per pot after seedling emergence. To control the effect of vernalization on flowering, trays were subjected to vernalization at 4°C with an 8h light period twelve days after sowing for a period of 46 days. Afterwards, plants were transplanted to bigger pots. Three plants of each genotype were grown under long day conditions in a greenhouse with 16h light duration and 20°C/16°C day/night. In parallel, three plants of each genotype were grown under short day conditions in a climate chamber (8h light, 20°C/16°C). Heading date (BBCH55) was recorded on the main tiller of each plant. Within each treatment, plants were grown in fully randomized design. Phenotypic data was manually curated and outlier test was performed. ANOVA, heritability and Best linear unbiased estimates (BLUEs) were calculated. Genotypes and their haplotypes considered for this study are indicted in Table S4.

#### *Heading date under vernalized and non-vernalized long-day field conditions at IPK*

Seeds were sown in turf trays at day 56 (vernalized treatment) and at day 99 (non-vernalized treatment) of the Julian calendar in 2010. Plants of the vernalized treatment received 43 days of vernalization in a climate chamber with 8h light and 4°C constant temperature. The experiment was conducted with two treatments (vernalized and non-vernalized) each with 2 replications. In each replication, 2 plants from each genotype were considered (in total 4 plants per genotype per treatment). Plants of both treatments (similar developmental stage) were transplanted to the field at day 119 of the Julian calendar (30^th^ of April) to avoid an effect of potential natural vernalization under field conditions in Germany. In the vernalized treatment, the following traits were scored: date of physiological maturity, growth habit (prostrate, intermedium and erect-type), plant height, flag leaf width, ear length and ear width. Outliers, repeatability and BLUEs were calculated as reported earlier.

### **Analysis of Environmental Data**

We used Principal Component Analysis (PCA) to rank the contribution of each bioclimatic variable. PCA was conducted using the R package FactoMiner (Le et al. 2008).

To infer environmental clusters of 1375 collection sites, Discriminant Analysis of Principal Components (DAPC) using the R package Adegent was employed (Jombart et al. 2010). The prior number of clusters were set to 2 to 20. Intra-correlated variables with high correlations exceeding r=0.9 were removed to allow spatial resolution. A total of 15 different indexes were used to test the possible number of clusters.

### **Supplementary Results**

### ***Genetic diversity at PPD-H1 within the Diversity panel***

Wild barleys from Israel possessed the highest genetic diversity (47 haplotypes), followed by Turkey (19 haplotypes). The number of haplotypes mostly decreases from Israel to the west: Israel (N=47), Cyprus (N=3), Greece (N=2) and Libya (N=1), and from Israel to the east: Israel (N=47), Jordan (N=9), Lebanon (N=4), Syria (N=8), Turkey (N=19), Iraq (N=2), Iran (N=5), Afghanistan (N=2), Turkmenistan (N=1), Uzbekistan (N=1) and Tajikistan (N=1). Several haplotypes were region-specific: for example, H34, H39 and H66 were specific to Israel.

Sixty-three haplotypes were unique to wild barley and not exploited in the domesticated barleys, which might harbor a key to local environment adaptation (Table S11). Interestingly, several of the non-exploited haplotypes clustered together, like (i) H48, H36, H38, H39, H85, H28, H5, H29, or (ii) H16, H62, H78, H34, H46 and H67 (Fig. 5). Moreover, these haplotypes were distinct and differed from the closest major haplotypes by several base pairs. For example, H48 differed from H7 by 10 SNPs or H62 by 9 SNPs from H4 (Fig. 5).

### ***Phylogenetic relationships and geographic distribution of photoperiod responsive and non-responsive haplotypes***

**Geographical group 1**: Also among the wild genotypes harboring haplotype H10 were three genotypes from Iran originating probably from two locations in the Khuzestan province. Accessions HOR2882 and IG112787 were collected by H. Kuckuck, Germany, during 1952-1954 and conserved at IPK and then shared with ICARDA. One genotype (PI249983) was collected by P.F. Knowles, University of California in 1958 and maintained at USDA.

Haplotype H10 was not found in Turkey despite extensively sampling of the entire distribution area of the species in the country (N=362). The disjunctive distribution of haplotype H10 suggests similar environmental conditions at the respective collection sites. It is interesting to note that a distinctive geographic distribution was also observed in few additional haplotypes which were collected in the western as well as in the eastern Fertile Crescent: e.g. H5 (Israel and Iraq) or H28 (Cyprus/Israel and Iran).

Interestingly, the non-responsive haplotypes were also found in domesticated barley from the Fertile Crescent: Haplotype H1 was found in 53 barley landraces, including one genotype from Israel (Bedouin landrace, newly collected by Hübner et al. (2009), Syria (N=1), Turkey (N=5) and Iran (N=10). Moreover, H1 was found in cultivars from Israel (N=2) and Iran (N=1). Haplotype H2, which is derived from H1, was detected in 56 landraces including four genotypes from Israel (incl. two Bedouin landraces, Hübner et al. 2009) and one genotype from Turkey. H2 was found in 13 cultivars from Israel and four from Turkey.

**Geographical group 2**: **H7->H9:** Haplotype H9 was carried by 34 landraces collected from Jordan, Syria, Turkey, Iraq, Iran and Yemen but also found in three wild barleys from Iran (from one location).

**H7->H45->H86:** Haplotype H45 proved to be region-specific for landraces from Afghanistan, Pakistan and India, and was found in three *H. agriocrithon* genotypes from China. Haplotype H86 was unique for one landrace from Afghanistan. Most genotypes carrying these haplotypes (H92 and H45) were classified as winter types (Table S4).

**Geographical group 3**: H75 -> H13 -> H4 -> H3 -> H95: Haplotype H13 was detected in (i) 13 wild barleys including 10 genotypes from Turkey, two from Syria and one from Afghanistan; and (ii) one landrace from Chad. **Haplotype H4** represents a major haplotype and was detected in 230 genotypes (135 = 14.33% of wild and 8.55% of domesticated). Wild barleys were collected in Greece (N=1), Israel (N=54, including the Desert type barley FT143), Jordan (N=6), Lebanon (N=4), Syria (N=6) and Turkey (N=64). The remaining were 39 landraces mainly from the Fertile Crescent but also from Libya (N=3), and 56 cultivars mainly from Turkey. All wild barleys harboring haplotype H3 (N=18) were collected west of Gaziantep in Turkey. Domesticated barley containing H3 were 28 landraces from 13 countries and 70 cultivars from 26 countries. The derived haplotype H95 was found in five cultivars from Europe.

**Geographical group 4**: H6 and derived haplotypes: Haplotype H6 was the most frequent haplotype and detected in 502 genotypes: 40.44% (N=381) of wild barley, 10.72% (N=119) of domesticated barley (40 landraces, 77 cultivars) and two of *H. agriocrithon*. The haplotype H6 was most frequent in wild barley and showed a more eastern distribution compared to the haplotype H7. The frequency of H6 increased from west to east: Israel (2.2%) - Jordan (11.1%) - Lebanon (12.5%) - Syria (54.1%) - Turkey (60.5%) - Iraq (93.1%) - Iran (90.1%) - Afghanistan (85.7%) - Turkmenistan (100%) - Uzbekistan (100%) - Tajikistan (100%). Several population/ region-specific haplotypes (all unique to Turkey) derived from H6, for example: H41, H49, H50 and H53.

### ***Analysis of environmental data shed more light on the region of origin of photoperiod non-responsiveness***

The collection sites of wild barley from Israel containing haplotype 10 are more similar to the Masada environment than the collection sites of haplotype 10 wild barley from Iran. DAPC analysis provided complementary results. The most important variables were Bio14, Bio15, Bio18, Bio9, Bio10, and Bio19, respectively (Fig. S11). Four environmental clusters were detected (Figs. S12, S13; Table S17b). Cluster 1 contained 543 wild barleys, mainly from the central and eastern part of the Fertile Crescent and 194 landraces. Among the wilds were the three wild barleys from Iran harboring H10. Cluster 4 contained 505 genotypes. A total of 397 wild barley collected mainly from the Eastern Mediterranean and the Near East were assigned to this cluster (including all 363 samples from Israel, and thus all samples containing H10 from Israel. In addition, 108 landraces, mainly from Africa, the Arabian Peninsula and the Near East, as well as the Masada barley were included in cluster 4 (Table S17). Photoperiod non-responsive barley was found in all four clusters.

The environmentally third closest collection site to Masada is the collection site of B1K-05 (Israel, Neomi), where FT013, FT014, FT015 and FT016 (all Desert type) were collected. All wild barleys from this population carried H66 at *PPD-H1*. FT013, FT015 and FT016 were also re-sequenced at *Btr1* and *Btr2* (Pourkheirandish et al. (2015) and were among the closest wild barleys to *btr1* (Fig. 7; Table S4).

### ***Vernalization requirement and phenotypic performance of genotypes containing haplotype H10 under long-day conditions***

In a field trial at IPK in 2010, under vernalized and non-vernalized conditions, heading date of 843 genotypes of wild and domesticated barley was investigated to determine their vernalization requirement and to characterize key agronomic traits in the vernalized treatment. For all traits, the repeatability of data was high and ranged from 0.86 to 0.98.

In the non-vernalized treatment, 582 genotypes were heading, while the remaining 261 genotypes (30%) were not flowering and therefore considered winter types.

On average, heading was reached 15 days later under non-vernalized compared to vernalized conditions among the set of 582 genotypes. Interestingly, a bimodal distribution of the heading date differences was observed, indicating two phenotypic groups - spring and facultative growth habit (Fig. S8a). In group 1, there were 232 genotypes with relatively similar heading dates in both treatments, where the range of heading date difference was -7 to +14 days (average heading date difference: +5.9 days). Group 2 consisted of 350 genotypes with a delayed heading date under non-vernalized conditions and the difference ranged from 14.2 to 35.8 days (average heading-time difference: +23.9 days). Genotypes from group 1 were classified as spring (flowering without vernalization), genotypes from group 2 as facultative types (flowering without vernalization but earlier when vernalized).

Of all 843 barley genotypes, 97 wild accessions did not exhibit completely wild characteristics and were excluded from subsequent analysis, leaving 746 accessions for comparison (470 wild and 276 domesticated). Based on our experiments that depict bimodal distribution, 204 accessions were classified as spring types (6 were wild barley from Israel) and 305 as facultative, while 237 accessions were classified as winter types. A total of 70 different haplotypes were represented within this panel. To compare the haplotypes for the phenotypic key characteristics assessed in the vernalized treatment, we excluded haplotypes present in less than five genotypes, resulting in a panel of 404 wild barley and 267 domesticated genotypes covering 17 different *PPD-H1* haplotypes.

The genotypes carrying the non-responsive haplotypes H1 and H2 at *PPD-H1* were mostly spring types. In contrast, the photoperiod responsive progenitor haplotype H10 was mainly found in facultative types (wild barley from Israel, N=9; and wild barley from Iran, N=3) but also in two spring types (wild barley FT147 from Israel, landrace FT537 from Turkey) and one winter type (wild barley FT002 from Israel) (Fig. S8b). The genotypes with haplotype H10 showed a short life cycle with the second earliest heading date (1. H66, 2. H10, 3. H26) and the earliest maturity date (1. H10, 2. H66, 3. H26) of all haplotypes under vernalized, long-day field conditions in Germany (Fig. S8c; Table S19), even if only wild barley was considered (Fig. S8d). In addition, the H10-containing genotypes were among the three genotypes with the shortest plant height, narrowest flag leaves, shortest main ear and narrowest main ear width (Table S19). From these data we conclude that plants containing haplotype 10 are well adapted to their local environmental conditions in the Southern Levant or in Khuzestan, and that they are characterized by facultative or even spring growth habit.

It should be noted that genotypes containing H10 or other haplotypes found in Desert-type wild barley (e.g. H26, H66) flowered later than haplotypes H1 and H2 under long-day and non-vernalized field conditions in Germany (Table S19). This is somewhat surprising because they flower (very) early and actually under short day conditions in their native environments in the Southern Levant but confirming their facultative growth habit. The only spring type wild barley with H10 (FT147 from Israel, Havarim stream) flowered very early in both treatments.

### **Supplementary Tables**

**Table S1:** List of 224 domesticated barley accessions (<https://gbis.ipk-gatersleben.de/>) considered to establish the GWAS panel. The 177 accessions are indicated, which were used for the GWAS analysis. More details are provided in Table S4.

**Table S2:** Details of multi-location field trials. The country, location, year, field-design and geographical coordinates are provided.

**Table S3:** List of 138 excluded genotypes which possessed non-wild or other uncertain characteristics.

**Table S4:** Overview of the geo-referenced Diversity panel. Passport data are provided for all 2057 genotypes. Haplotype numbers are provided for all re-sequenced fragments. Assigned haplotype numbers for the 1367 bp and 898 bp fragments differed in three cases: haplotypes H4, H6 and H7 originally defined for the 898 bp fragment were divided into H4a/H4b, as H6a/H6b, and H7a/H7b for the 1367 bp fragment, respectively.

**Table S5:** Summary statistics of heading date (Hd) data from multi-location field trials (GWAS panel).

**Table S6:** Pearson correlation coefficient of the days to heading (Hd) from multi-locations field trials.

**Table S7:** List of significant SNPs from the GWAS analysis. Highlighted in Yellow are the SNPs from the *PpdH1* region. SNPs, chromosome, physical positions, significances, minus log10P-values, effects on the transcript (based on the Bayer et al. 2017).

**Table S8:** Analysis of variance (ANOVA) of the SNP x Site.

**Table S9:** Estimated SNP x Site effects.

**Table S10:** GWAS panel: haplotype frequency distribution according to the geographic origin of GWAS panel. The 1367 bp fragment was re-sequenced (Fig. 3; Fig. S2). The % of genotypes from the respective geographical groups are shown.

**Table S11:** List of haplotypes and their frequencies found in the Diversity panel.

**Table S12:** NCBI accession numbers corresponding to the 14 haplotypes found in the GWAS panel (1367 bp fragment).

**Table S13:** NCBI accession numbers corresponding to the 90 haplotypes found in the Diversity panel (898 bp fragment).

**Table S14:** Population statistics of 2057 re-sequenced genotypes at *PPD-H1* using the primer combination PP05 + PP04 that covered the CCT domain including SNP22 (causal SNP reported by Turner et al. 2005).

**Table S15:** Allelic states at *PPD-H1* and *HvCEN* of the 6000 years old barley sample JK3014.

**Table S16:** Values of the first five Principal Components (PCs) scores obtained from PCA of 19 bioclimatic variables of 1375 collection sites of wild and landrace barleys, and the ancient Masada barley.

**Table S17:** Values of 19 bioclimatic variables for 1375 sampling sites of wild barley, landraces and the ancient barley sample JK3014 used for PCA analysis. The values of Linear Discriminant scores (LD1 to LD3), and the inferred environmental cluster of each barley obtained from DAPC analysis are shown.

**Table S18:** Overview of closest current environmental conditions at collection sites of extant wild and landrace barley compared to the collection site of the ancient Masada barley obtained from pairwise Euclidean distances based on bioclimatic variables. K1: closest H10 containing genotypes to the Masada sample; K2: closest wild barleys from Israel to the Masada sample; K3: closest wild barleys from all countries to the Masada sample; K4: closest landraces from Fertile Crescent carrying H1 to the Masada sample; K5: closest landraces and cultivars from the Fertile Crescent (countries: CYP, EGY, IRN, TUR, ISR, SYR, LBN, JOR, IRQ) to the Masada sample; K6: closest wild barleys and all samples considered in K5 to the Masada sample. See Tables S4, S17 for more information.

**Table S19:** Evaluation of vernalization requirement and key agronomic traits in a panel of 671 domesticated and wild barley (corresponding to 17 haplotypes of *PPD-H1* with at least 5 genotypes per haplotype) evaluated in the field in Gatersleben, Germany in 2010. Repeatability of phenotypic traits is presented along with the average values of haplotypes based on phenotypic BLUEs.

### **Supplementary Figures**

**Fig. S1:** Heading date variation across the three locations where *PPD-H1* was QTL based on the putative casual SNPs. Box plots displaying days to heading (Hd) considering SNP22 (T/G) and SNP48 (T/C).

**Fig. S2:** Phylogenetic networks derived from re-sequenced *PPD-H1* fragments (1367 bp) based on the GWAS panel. (a) MJ-network (as in Fig. 5) but based on the 14 haplotypes identified in the GWAS panel (Table S4). (b) NeighborNet network computed for these 14 haplotypes.

**Fig. S3:** Differential response of photoperiod responsive and non-responsive genotypes of the GWAS panel during four developmental stages under inductive long day conditions in the greenhouse at IPK. Lower Growing Degree Days (GDD) values for haplotype H10 containing genotypes (BCC533, BCC759) compared to photoperiod non-responsive genotypes indicate accelerated heading dates for haplotype H10 containing genotypes (Alqudah et al. 2014). This provides further evidence that H10 containing genotypes are photoperiod responsive.

Thermal time measured as GDD from sowing to awn-primordium, tipping, heading and anther extrusion stages is provided (y-axis). Blue: photoperiod non-responsive haplotype (*ppd-H1*) containing genotypes; Grey: photoperiod responsive haplotype (*Ppd-H1*) containing genotypes; Orange: H10 containing genotypes. See Figs. 3b, 5; Fig. S2; Table S4.

**Fig. S4:** Geographical distribution of the Diversity panel. (a) Overview of the distribution of the 2057 accessions on the world-map (red = *H. spontaneum*; yellow = *H.* *agriocrithon* and blue = *H. vulgare*). The Fertile-Crescent region is enlarged; (b) geographical distribution of photoperiod non-responsive, and (c) geographical distribution of photoperiod responsive haplotypes.

**Fig. S5:** Polymorphic sites defining the 90 haplotypes detected within 2057 re-sequenced genotypes of the Diversity panel. SNP22 is indicated by a red rectangular frame (central position, T/G SNP). Red dot on the left side, haplotype found in wild barley; dark blue dot, haplotype found in domesticated barley; dark blue and red dots, haplotype found in domesticated and wild barley.

**Fig. S6:** Geographical distribution of haplotype H10 containing genotypes (red = wild barley; blue = domesticated barley). In total, haplotype H10 was found in 16 wild barleys and 4 domesticated barleys.

**Fig. S7:** Geographical clines of haplotypes at *PPD-H1* presented as boxplots. Comparison of latitude, longitude and altitude between (a) genotypes containing non-responsive haplotypes (H1 and H2), and (b) photoperiod non-responsive *vs.* responsive genotypes.

**Fig. S8:** Evaluation of vernalization requirement and phenology for 843 domesticated and wild barley genotypes of the Diversity panel grown under vernalized and non-vernalized field conditions at IPK. **(a)** Histogram of heading date differences between non-vernalized and vernalized conditions (heading date in non-vernalized minus heading date in vernalized conditions) for 585 genotypes that were flowering in both treatments. The rest of the panel was classified as winter type (not flowered under non-vernalized conditions). **(b)** Overview of the frequency of the classified growth habit type among 17 *PPD-H1* haplotypes carried by at least 5 genotypes (in total 671 genotypes). The numbers on top refer to the number of genotypes that carry the corresponding haplotype. **(c)** Time to heading and to maturity in the vernalized treatment in wild and domesticated barley for 17 *PPD-H1* haplotypes carried by at least 5 genotypes (in total 671 genotypes. **(d)** Time to heading and to maturity in the vernalized treatment in wild barley for 11 *PPD-H1* haplotypes carried by at least 5 genotypes (in total 396 genotypes). See Table S3 and Table S4 for more information.

**Fig. S9:** Visualization of the first five principal component scores and their relationship with the original bioclimatic variables. A bolded vector indicates a strong positive (green) or negative (red) correlation of the original bioclimatic variable to the respective PC score.

**Fig. S10:** Box plots of geographical and bioclimatic variables for the environmental conditions at collection sites of major *PPD-H1* haplotypes (H1, H2, H3, H4, H6, H7) and haplotypes H10 and H75, for wild and landrace barley (Lon, Lat, Alt, Bio1-Bio19). The vertical bar inside the box plot is the median value and the red dot indicates the mean value. H01.FC: H1 containing photoperiod non-responsive barley from the Fertile Crescent; H02.AF: H2 containing photoperiod non-responsive barley from Africa; H10.IRN: H10 containing wild barley from Iran; H10.ISR: H10 containing wild barley from Israel. Landraces: Landraces with remaining haplotypes; Wild: wild barley with remaining haplotypes.

**Fig. S11:** Contribution of bioclimatic variables to the first two discriminant functions applying the threshold loading of 0.05. The most important bioclimatic variables were Bio14, Bio18, Bio15, Bio9, Bio10, and Bio19, respectively.

**Fig. S12:** DAPC biplot. Graphical representation for the first two discriminant functions. The plot indicates the separation of 1375 samples into four clusters.

**Fig. S13:** Density plot of individuals along first discriminant function with different colors from the discriminant analysis of principal components (DAPC). The four clusters are shown.
